# Full Spectrum Flow Cytometry for High-Dimensional Immunophenotyping of Mouse Innate Lymphoid Cells

**DOI:** 10.1101/2022.01.05.475054

**Authors:** Kyle T. Mincham, Robert J. Snelgrove

**Affiliations:** National Heart and Lung Institute, Imperial College London, London, SW7 2AZ

**Author notes:** Corresponding author. Address: National Heart and Lung Institute, Sir Alexander Fleming Building, Imperial College London, South Kensington, London, SW7 2AZ.

## Abstract

This 25-parameter, 22-colour full spectrum flow cytometry panel was designed and optimised for the comprehensive enumeration and functional characterisation of innate lymphoid cell (ILC) subsets in mouse tissues (Table 1). The panel presented here allows the discrimination of ILC progenitors (ILCP), ILC1, ILC2, NCR^+^ ILC3, NCR^−^ ILC3, CCR6^+^ lymphoid tissue-inducer (LTi)-like ILC3 and mature natural killer (NK) cell populations. Further characterisation of ILC and NK cell functional profiles in response to stimulation is provided by the inclusion of subset-specific cytokine markers, and proliferation markers. Development and optimisation of this panel was performed on freshly isolated cells from adult BALB/c lungs and small intestine lamina propria, and *ex vivo* stimulation with phorbol 12-myrisate 13-acetate, ionomycin and pro-ILC activating cytokines.

**Ethical compliance statement:** All mouse experiments were performed in accordance with the recommendations in the Guide for the Use of Laboratory Animals of Imperial College London, with the ARRIVE (Animal Research: Reporting of In Vivo Experiments) guidelines. All animal procedures and care conformed strictly to the UK Home Office Guidelines under the Animals (Scientific Procedures) Act 1986, and the protocols were approved by the Home Office of Great Britain.

## Background

Innate lymphoid cells (ILCs) are a unique subset of innate effector cells enriched at mucosal surfaces, with diverse roles in host defence, tissue remodelling and repair, inflammation and metabolic homeostasis^1^. Despite lacking rearranged antigen receptors, ILCs display remarkable homology with conventional T helper (Th) type 1 (Th1), Th2 and Th17 cells in regards to phenotype and function, and thus are similarly classified into ILC1, ILC2 and ILC3 subsets^2^. Moreover, *bona fide* ILC subsets have now been expanded to include both cytotoxic natural killer (NK) and lymphoid tissue-inducer (LTi) cells, which are phenotypically and functionally similar to ILC1 and ILC3 respectively, yet exhibit distinct developmental trajectories^3^. However, it is increasingly recognised that ILC subsets are not fixed, and that these cells can exhibit significant plasticity depending on the local inflammatory milieu^4–8^. As such, ILCs demonstrate intra-subset phenotypic heterogeneity depending on their specific microenvironment^9, 10^, and deep immunophenotyping therefore requires a comprehensive array of phenotypic and functional markers to accurately capture this biological variation.

Although crucial in multiple biological settings, ILCs constitute a relatively minor population within both mouse and human lymphoid and non-lymphoid tissues and blood, comprising 1-5% of CD45^+^ leukocytes^5, 11–15^. Due to this inherent scarcity, acquiring as much information as possible on a single-cell level is of upmost importance for accurate ILC discrimination. High resolution ILC characterisation is further convoluted in tissues such as the lung and small intestine, where the intrinsically autofluorescent nature of the samples results in a heightened background noise-to-signal ratio. With this in mind, cellular characterisation via full spectrum flow cytometry provides a technological advancement given its high parameter capabilities^16^ combined with the capacity to extract cellular autofluorescence profiles, thus improving discrimination of rare populations compared to conventional flow cytometry^17^. As such, the full spectrum panel described herein enables the detailed enumeration and functional assessment of all ILC subsets identified within mouse tissues, with a specific focus on the characterisation of lung and small intestine lamina propria (siLP) populations given the divergent repertoire of resident ILC subsets localised to these tissue compartments.

ILCs arise from common lymphoid progenitors within the bone marrow (BM), which through a series of intermediate commitment stages give rise to ILC progenitors (ILCP). While initially believed to be restricted within the BM, ILCPs are now recognised to exist within circulation and as tissue-resident populations at distal sites in both neonates and adults, including the lungs, skin, and secondary lymphoid tissues^18–21^. During early development, expression of the transcription factor promyelocytic leukaemia zinc finger (PLZF; encoded by *Zbtb16*) dictates the bifurcation of innate lymphoid progenitors, whereby PLZF^+^ ILCPs are restricted to the generation of ILC1, ILC2 and ILC3, but not LTi or NK cells^22, 23^. In addition to PLZF, ILCPs display high-level expression of *Il18r1*, and thus a requirement for IL-18 signalling through IL-18Rα for proliferation and differentiation^19^. Importantly, both ILCPs and mature ILC subset are dependent on canonical IL-7 signalling for development and function, with expression of the receptor (IL-7Rα) and consumption of IL-7 dramatically greater in ILCs than their T cell counterparts^24^. Continual differentiation of IL-7Rα^+^PLZF^+^IL-18Rα^+^ ILCP into distinct ILC subsets is under the control of a series of lineage-defining master transcription factors, with all terminal subsets lacking expression of extracellular markers routinely used to identify both lymphoid and myeloid cells, subsequently defining ILCs as lineage negative (Lin^−^) cells (refer to Table 2 for all appropriate lineage markers).

**Table 1.** Summary Table

|  |  |
| --- | --- |
| Purpose | Deep immunophenotyping and functional assessment of ILC subsets |
| Species | Mouse |
| Cell types | Innate Lymphoid Cells |
| Cross-reference | N/A |

**Table 2.** Reagents used for OMIP

| Specificity | Fluorochrome | Clone | Purpose |
| --- | --- | --- | --- |
| CD45 | BUV395 | 30-F11 | Pan leukocytes |
| IL-7Rα | PE-Cy5 | SB/199 | Pan ILC |
| PLZF | PE | MAGS21F7 | ILCP |
| IL-18Rα | PerCP-eFluor 710 | P3TUNYA | ILCP and ILC1 |
| T-bet | BV421 | 4B10 | ILC1 and ILC3 subsets |
| NKp46 | PE/Dazzle 594 | 29A1.4 | ILC1, ILC3 subsets and NK cells |
| CD49a | BUV496 | Ha31/8 | ILC1 subsets and trNK cells |
| TRAIL | PE-Cy7 | N2B2 | ILC1 subsets |
| IFN-γ | BUV737 | XMG1.2 | ILC1 and NK cells |
| GATA-3 | BB700 | L50-823 | ILC2 |
| KLRG1 | BV480 | 2F1 | ILC2 |
| ST2 | R718 | U29-93 | ILC2 |
| IL-5 | APC | TRFK5 | ILC2 |
| IL-13 | eFluor 450 | eBio13A | ILC2 |
| RORγt | BV786 | Q31-378 | ILC3 |
| CCR6 | BV711 | 140706 | ILC3 subsets |
| IL-17A | BV650 | TC11-18H10.1 | ILC3 |
| IL-22 | Alexa Fluor 647 | Poly5164 | ILC3 |
| CD49b | eFluor 506 | DX5 | NK cells |
| Ki-67 | Alexa Fluor 532 | SolA15 | Proliferating cells |
| B220 | FITC | RA3-6B2 | B cell subsets (Lin dump) |
| CD3 | FITC | 145-2C11 | T cell subsets (Lin dump) |
| CD4 | FITC | GK1.5 | T cell subsets (Lin dump) |
| CD5 | FITC | 53-7.3 | T cell and B cell subsets (Lin dump) |
| CD11b | FITC | M1/70 | Myeloid/NK/Granulocytes (Lin dump) |
| CD11c | FITC | N418 | Myeloid cells/Granulocytes (Lin dump) |
| CD19 | FITC | 1D3/CD19 | B cell subsets (Lin dump) |
| F4/80 | FITC | BM8 | Macrophages (Lin dump) |
| FcεR1α | FITC | MAR-1 | Mast cells and Basophils (Lin dump) |
| Gr-1 | FITC | RB6-85C | Myeloid cells/Granulocytes (Lin dump) |
| TCRβ | FITC | H57-597 | $\alpha\beta$ T-cells (Lin dump) |
| TCR$\gamma\delta$ | FITC | UC7-13D5 | $\gamma\delta$ T-cells (Lin dump) |
| Ter119 | FITC | TER-119 | Red blood cells (Lin dump) |
| Dead cells | LIVE/DEAD™ Blue | - | Viable cells |

ILC1s are characterised by their constitutive expression of T-bet (encoded by *Tbx21*), which is central for their production of interferon (IFN)-γ and ensuing response to intracellular pathogens following IL-12 and IL-18 stimulation^18, 25^. Owing to their similarities with cytotoxic NK cells, ILC1s express NKp46, however tissue-specific distinctions can be made between these two subsets via the preferential expression of CD49a and/or TRAIL by ILC1, in conjunction with the absence of CD11b and CD49b^3, 18, 25^.

ILC2s represent the most abundant subset of ILCs in mouse lungs^5^ and are dependent on the expression of GATA-3 for maintenance and survival^26, 27^. Analogous to their Th2 counterparts, ILC2s play a key role in controlling helminth infection^26, 28, 29^, whilst perpetuating allergen-induced allergic inflammation^30–33^ via the production of IL-5 and IL-13 in response to alarmins IL-33, IL-25 and thymic stromal lymphopoietin (TSLP). Given their extensive roles in both immunity and inflammation, ILC2s are known to display dynamic expression of extracellular markers associated with pro-inflammatory functions, exemplified via variations in IL-33 receptor (ST2) and KLRG1 expression depending on the inflammatory signal initiating the primary response^13, 34, 35^. Moreover, the inflammation-dependent plasticity of ILC2s has been demonstrated in both mouse lungs and human peripheral blood, whereby stimulation with IL-1β, IL-12 and IL-18 promotes ILC2s to adopt an ILC1-like transcriptional and functional profile associated with the expression of T-bet (*Tbx21*) and production of IFN-γ^5, 6, 34, 36^. Similarly, human ILC2s, in response to IL-1β and IL-23 stimulation, transdifferentiate into a subset exhibiting ILC3-like characteristics, evidenced by upregulation of the ILC3 signature transcription factor RORγt and production of IL-17^8, 37^.

ILC3s are a heterogeneous subset and contribute broad roles in combating against extracellular microbes, including fungi and bacteria, via the production of IL-17A and IL-22 following IL-1β and IL-23-mediated activation^38–40^. Representing the dominant ILC subset within the steady state siLP^41^, ILC3s are strictly reliant upon RORγt (encoded by *Rorc*) expression for development and function^39, 42, 43^. However, mouse ILC3s can be further classified into 3 subtypes based on the differential expression of NKp46 and CCR6; namely NKp46^+^CCR6^−^ (NCR^+^), NKp46^−^CCR6^−^ (NCR^−^) and NKp46^−^CCR6^+^ (CCR6^+^ LTi-like) ILC3s^42, 44, 45^. Moreover, NCR^+^ ILC3s share transcriptional expression of T-bet with ILC1s, crucial for their expression of NKp46 and endowing them with the ability to produce IFN-γ^46, 47^. Furthermore, stimulation of mouse tissue-specific NCR^+^ ILC3s with IL-12 promotes downregulation of RORγt and concomitant upregulation of T-bet, further promoting a phenotype associated with IFN-γ^+^ ILC1s^4^ and demonstrating their plasticity in response to environmental cues.

Given this microenvironmental-driven impact on ILC heterogeneity, the developmental phase of this full spectrum flow cytometry panel involved the prioritisation of definitive ILCP, ILC1, ILC2 and ILC3 transcription factors, as reflected in the gating strategy (Figure 1). An added benefit of full spectrum flow cytometry is the ability to remove red blood cell contamination by plotting side-scatter (SSC) captured by the 405nm violet laser against SSC-B from the 488nm blue laser^48^ (Figure 1A), thus ensuring purity of downstream terminal cell populations. Viable leukocytes (LIVE/DEAD Blue^−^ CD45^+^; Figure 1A) were selected and total ILCs were defined by their expression of IL-7Rα^+^ and lack of lineage markers used to routinely define T cells, B cells, myeloid cells, and red blood cells (CD3, CD4, CD5, CD11b, CD11c, CD19, B220, F4/80, FcεR1, Gr1, TCRβ, TCRγδ, Ter119; Figure 1B). ILCPs within the lung were characterised by their ubiquitous expression of PLZF, in combination with IL-18Rα^+^ (Figure 1C). Mature ILC subsets within the lung (Figure 1D) and siLP (Figure 1E) were stratified as T-bet^+^RORγt^−^ST2^−^ CD49b^−^IL-18Rα^+^NKp46^+^CD49a^+/−^TRAIL^+/−^ ILC1s, GATA-3^+^T-bet^−^RORγt^−^KLRG1^+/−^ST2^+/−^ ILC2s and either RORγt^+^T-bet^+/−^NKp46^+^CCR6^−^ (NCR^+^), RORγt^+^T-bet^+/−^NKp46^−^CCR6^−^ (NCR^−^) or RORγt^+^T-bet^+/−^NKp46^−^CCR6^+^ (CCR6^+^ LTi-like) ILC3s. Regarding ILC2s, it is important to note that ST2 expression on *ex vivo* stimulated siLP ILC2s is largely absent (Figure 1E)^9, 49^. Moreover, while lung ILCs lack CD4 expression^50^, intestinal CCR6^+^ LTi-like ILC3s can be sub-classified as CD4^+^ and CD4^−51^. As such, the inclusion of CD4 within the lineage cocktail may influence the characterisation of ILC3 subsets on a tissue-specific basis. ILC cytokine production and proliferation within the lung (Figure 1F) and siLP (Figure 1G) was assessed in response to subset-specific activation signals via *ex vivo* stimulation with phorbol 12-myrisate 13-acetate (PMA), Ionomycin and Brefeldin A, and either IL-12 and IL-18 (pro-ILC1), IL-33 (pro-ILC2) or IL-1β and IL-23 (pro-ILC3). Stimulation with ILC subset-specific cytokines clearly demonstrates a heightened propensity for IL-13 and IL-22 production by siLP ILC2s and ILC3 subsets respectively, whereas lung ILC1s significantly upregulate IFN-γ compared to their siLP counterparts. Although the focus of this panel was to accurately define ILCPs and ILC1-3 subsets, inclusion of CD11b within the lineage cocktail enables the enumeration of conventional mature CD11b^+^NKp46^+^T-bet^+^ NK cells (Figure 1H)^52^, with additional characterisation of lung tissue-resident (trNK) and circulating (cNK) populations on the basis of CD49a and CD49b expression^53^ (Figure 1H). Markers including KLRG1^54^ and RORγt^55^, and cytokines IFN-γ, IL-5, IL-13^56^ and IL-22^57^ are all reportedly expressed by NK cell subsets based on maturation, tissue localisation and disease state, further highlighting the diversity of this OMIP.

**Figure 1.**
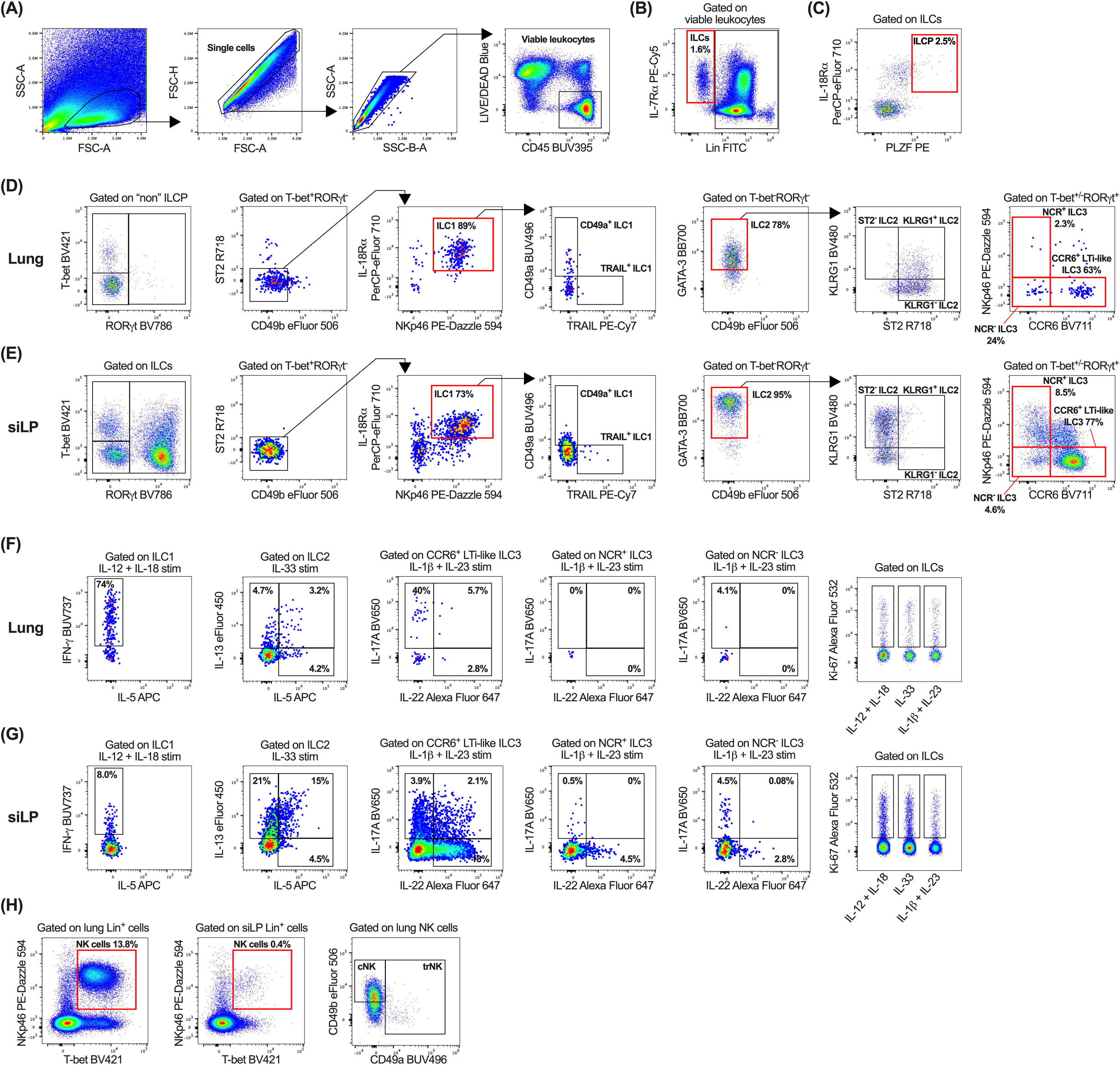
Gating strategy to characterise ILC populations and their subset-specific cytokine and proliferative profiles. After exclusion of cellular debris, doublets, red blood cell contamination and non-viable CD45^+/−^ cells, viable CD45^+^ leukocytes **(A)** were gated on IL-7Rα and lineage (CD3, CD4, CD5, CD11b, CD11c, CD19, B220, F4/80, FcεR1, Gr1, TCRβ, TCRγδ, Ter119; Lin) negative cells to identify ILCs **(B)**. ILCPs were then identified as IL-18Rα^+^PLZF^+^ cells **(C)**. Characterisation of ILC1, ILC2, NCR^+^, NCR^−^ and CCR6^+^ LTi-like ILC3 subsets in the lung **(D)** and siLP **(E)**. All samples in panel A-E were *ex vivo* stimulated for 5 hours with PMA/Ionomycin/BFA. Single cell suspensions were *ex vivo* stimulated for 5 hours with PMA/Ionomycin/BFA and either IL-12 and IL-18 (pro-ILC1), IL-33 (pro-ILC2) or IL-1β and IL-23 (pro-ILC3) and ILC subsets were assessed for production of subset-specific cytokines (IFN-γ, IL-5, IL-13, IL-17A and IL-22) and cellular proliferation (Ki-67) within the lung **(F)** and siLP **(G)**. **(H)** Identification of NK cells within the lung and siLP, with downstream characterisation of lung tissue-resident NK (trNK) and circulating NK (cNK) cells. Plots in panel A-C are representative of individual lung samples. For panels B-H, viable leukocytes from 3 technical lung or siLP replicates were concatenated into single FCS files prior to gating. All fluorochrome-conjugated antibodies were titrated (Online Figure 1) during panel optimisation and manual gating was determined using fluorescence minus one controls (Online Figure 2) where necessary.

In summary, we present here the first full spectrum flow cytometry panel (Table 2) to comprehensively profile the phenotypic and functional state of ILCs in multiple mouse tissues. Moreover, by maintaining availability on the 355nm ultraviolet laser, this panel can be expanded to incorporate additional markers of interest with little impact on the overall Complexity™ Index (a value of how distinct a collection of spectral signatures are from one other when unmixed simultaneously), thereby providing a valuable resource within the rapidly expanding field of ILC biology.

## Acknowledgement

The authors would like to thank Dr James Harker and the Imperial College London Flow Cytometry Facility South Kensington for their valuable advice on Cytek^®^ Aurora operation during the initial design of this panel. In addition, the authors would like to thank Christopher McRandle and Dr Adam Davison from Cytek^®^ Biosciences for their advice on data quality control using SpectroFlo^®^ software. The authors acknowledge financial support from Imperial College London through an Imperial College Research Fellowship grant awarded to K.T.M. R.J.S. is a Wellcome Trust Senior Research Fellow in Basic Biomedical Sciences (209458/Z/17/Z). Parts of this work were also funded through a Rosetrees Trust/The Stoneygate Trust Project Grant (PGS21/10072).

