## Supplementary material for "Full Spectrum Flow Cytometry for High-Dimensional Immunophenotyping of Mouse Innate Lymphoid Cells"

### Panel Development Strategy

#### *Instrument configuration*

This 25-parameter, 22-colour full spectrum flow cytometry panel was developed on a 5 laser 64 detector Cytek® Aurora (Cytek Biosciences) as per the optical configuration detailed in Online Table 1. Gain settings were calibrated daily using SpectroFlo® QC Beads.

#### *Antibody-fluorochrome titrations*

All antibodies used in the initial development phase and final optimisation of this OMIP were titrated using serial 2-fold dilutions to determine the optimal concentration for use (Online Figure 1). All titrations were performed on cells that had undergone *ex vivo* stimulation with PMA, ionomycin and brefeldin A for 5 hours, followed by fixation and permeabilisation regardless of intracellular staining, as this is the protocol used for the fully-stained samples. Optimal titrations were chosen based on visual inspection of positive and negative population separation rather than greatest saturation (highest signal-to-noise ratio whilst still achieving saturation) in order to reduce fluorochrome spreading and background signal. As seen in Online Figures 1 and 2, fluorochrome sensitivity using full spectrum flow cytometry is extremely high, and simply using titrations optimised on a conventional cytometer (Online Figure 3) will not suffice.

#### *Antibody and fluorochrome selection*

The primary stage of panel development required the definition of a general gating strategy (for final version see Figure 1), as this is an ILC-specific panel and therefore many of the markers necessary for accurate characterisation are co-expressed. Accurately defining co-expressed markers aids in the appropriate distribution of fluorochromes, thereby avoiding issues associated with fluorochrome spillover and subsequent loss of marker resolution. The assignment of antigen-fluorochrome combinations was then based around antigen density, Cytek® Aurora fluorochrome sensitivity/brightness (located at cytekbio.com/pages/fluorochrome-guides) and commercial availability, with all iterations and the final OMIP detailed within Online Table 2 and 3 respectively.

Firstly, to ensure all components of the lineage cocktail could be purchased on the same fluorochrome, and to maximise panel cost effectiveness (given there are 13 lineage markers within the panel) without sacrificing population resolution, all lineage markers were assigned to FITC. CD45 was then allocated to APC-Cy7 due to its constitutively high antigen density, in combination with Fixable Viability Stain 575V (FVS 575V) for the identification of viable leukocytes, based on successful panels previously developed on a conventional cytometer[1]. Due to the rarity of ILCs, maximal use of bright fluorochromes was important for adequate cellular resolution. Priority was therefore assigned to the constitutively expressed IL-7Rα and core transcription factors essential to ILCP (PLZF), ILC1 (T-bet), ILC2 (GATA-3) and ILC3 (RORγt) characterisation. IL-7Rα was assigned to PE-Cy5 due to both its brightness and clear separation from the FITC peak in the B2 emission channel (Online Figure 4), ensuring clear distinction of IL-7Rα$^{+}$Lin$^{-}$ ILCs from the large and heterogeneous Lin$^{+}$ population present within all tissues assessed (Figure 1 and Online Figure 5). To avoid spillover and associated signal loss of the core transcription factors, their assigned fluorochromes were distributed over the ultraviolet (UV), violet (V) and yellow-green (YG) lasers. An additional consideration for T-bet and RORγt was their context specific co-expression on ILC3 subsets, therefore BV421 was designated to T-bet and BV786 to ROR$\gamma$t, as these fluorochromes demonstrate considerable brightness and the 405nm violet laser provides tight emission bands. Due to limited commercial availability on bright fluorochromes, PLZF was assigned to PE. ILC2s are the dominant subset within the lungs[2], and therefore the moderately bright BUV395 was selected for GATA-3, which has limited spillover across the entire spectrum and allows the reservation of brighter fluorochromes for downstream ILC2 subset markers.

For ILC1 characterisation, PerCP-eFluor 710 and PE/Dazzle 594 were selected for IL-18R$\alpha$ and NKp46 (as opposed to NK1.1 so as to not constrain this OMIP to a specific mouse strain) respectively. This combination of very bright fluorochromes will provide a distinct double-positive population of ILC1s, clearly defined from CD49b eFluor 506 on the violet laser, which allows for the accurate removal of NK cell contamination within the ILC1 population. To account for tissue-specific variations in CD49a and TRAIL expression by ILC1s[3], and uncertainties regarding their co-expression, these markers were assigned to opposing ends of the available spectrum. Commercial availability of anti-mouse TRAIL (CD253) is limited and was subsequently designated to PE-Cy7, while BUV496 was reserved for CD49a as its proximity to GATA-3 BUV395 will not influence signal resolution due to an absence of co-expression. It is important to note that the BUV496 UV7 channel exhibits a high degree of autofluorescence in mouse lungs (Online Figure 6). However, this autofluorescent signature can be accounted for during data quality control using SpectroFlo® and the appropriate autofluorescence extraction method. Lastly, BUV737 was selected for IFN-$\gamma$ due to its high fluorescence intensity, which is dramatically upregulated on both ILC1s (Online Figure 7) and NCR$^+$ ILC3s (Online Figure 8) following stimulation with IL-12 and IL-18.

ILC2 markers KLRG1 and ST2 can be broadly classified as tertiary markers due to their variable tissue- and microenvironmental-specific expression. Such markers benefit from conjugation to a bright fluorochrome and were assigned to BV480 and R718 (a brighter alternative to APC-R700) respectively. It needs to be noted that the BUV737 (IFN-$\gamma$) acceptor dye can be excited by the 640nm red laser, resulting in low-level emission in the 700nm+ range. However, ST2 and IFN-$\gamma$ are not co-expressed and therefore spillover should be monitored but minimally disruptive. Regarding the characterisation of tissue-specific ILC subsets, it is noteworthy that phenotypic marker expression profiles remain consistent on all subsets irrespective of stimulation condition (Online Figures 9-11), with the exception of ST2 which appears to be downregulated on ILC2s following PMA/Ionomycin stimulation (Online Figures 10 and 11). ILC2-specific cytokines IL-5 and IL-13 were then allocated to APC and eFluor 450 respectively. While the bright APC signal will not be impacted by spread from other fluorochromes within the panel, the close emission proximity of IL-13 eFluor 450 in the V3 channel with T-bet BV421 in the V1 channel needs to be observed for spillover. However, as above, this issue is largely mitigated by a lack of co-expression.

Given the classification of ILC3 subsets on the basis of NKp46 or CCR6 expression, BV711 was assigned to CCR6, ensuring appropriate subset separation with negligible spillover into the parent ROR$\gamma$t BV786 population. Due to limited availability, IL-22 PerCP-Cy5.5 was selected based on its high stability in fixative and recommendation for intracellular staining. BV650 was then selected for IL-17A due to its tight emission band, high degree of brightness, and negligible spillover across the entire spectrum.

Finally, Ki-67 was assigned to Alexa Fluor 532, as any potential spillover from FITC is mitigated by the selection of lineage (FITC) negative cells during early ILC identification. Visual inspection of Ki-67 expression on each discrete ILC subset (Online Figure 12) additionally confirmed the Alexa Fluor 532 signal was not being captured within the $Lin^+$ FITC gate, thus validating Ki-67 fluorochrome selection was within the optimal region of the spectrum for terminal selection gates.

***Panel optimisation***

*In silico* optimisation was performed using the Similarity Index Matrix (SIM; Cytek Biosciences), with a similarity index (SI) between 2 fluorochromes ≤0.98 indicating their spectral signature is different enough to be used in combination[4]. From a total of 231 fluorochrome combinations, only 3 had a SI ≥ 0.8; eFluor450 and BV421 (SI = 0.85), eFluor506 and BV480 (SI = 0.87) and PerCP-eFluor 710 and PerCP-Cy5.5 (SI = 0.86; data not shown but available from https://spectrum.cytekbio.com/). While these SI values indicate the spectral profiles are distinct enough to be used in combination, consideration was taken during panel design to ensure markers used on these fluorochromes were not co-expressed.

Following computational optimisation, fluorochrome spreading was assessed using both cell and bead single stain reference controls to generate a Spillover Spreading Matrix (SSM; Online Table 4) using FlowJo™ (BD Biosciences; version 10.8.1). The initial SSM identified several problematic fluorochrome combinations that may result in loss of signal resolution in the secondary detectors, notably:

- BV421 into BV480: Co-expression does not occur between T-bet (BV421) and KLRG1 (BV480) and therefore BV421 will not impact signal resolution of BV480. This was confirmed upon visual inspection.

- BV786 into FITC: While co-expression will be present between RORγt (BV786) and lineage (FITC) positive cells, the first delineation gate for ILCs within the gating strategy is the exclusion of $FITC^+$ lineage cells, therefore minimising any potential spread issues, as confirmed upon visual inspection

- PerCP-eFluor 710 into R718: Co-expression is largely non-existent between IL-18Rα (PerCP-eFluor 710) and ST2 (R718), as evident in Figure 1C. Visual inspection determined this spread does not impact positive signal resolution of ST2.

- PE-Cy5 and PerCP-eFluor 710 into PerCP-Cy5.5: Although spectral similarities and potential spreading between PerCP-eFluor 710 and PerCP-Cy5.5 were also identified by the SIM, these markers are not co-expressed. However, all ILC subsets will be PE-Cy5 (IL-7Rα) positive. Due to the scarcity of IL-$22^+$ ILC3s within the lungs, and the relatively dim signal of the PerCP-Cy5.5 fluorochrome, spreading from PE-Cy5 posed a significant issue with the current iteration of the panel that needed to be addressed.

- PE-Cy5, PE/Dazzle 594 and PerCP-eFluor 710 into FVS 575V: All indicated fluorochromes display a spreading value >7 into the FVS 575V channel. Visual inspection demonstrated abnormal spreading within the viable (FVS $575V^-$) population for both PE-Cy5 and PE/Dazzle 594 (Online Figure 13A). It was therefore decided to move the viability stain to LIVE/DEAD™ Fixable Blue (Online Figure 13B) due to its unique signature, while additionally freeing up the V10 channel for future panel expansion.

- R718 into APC-Cy7: Visual inspection of the spread occurring between R718 (ST2) and APC-Cy7 (CD45) revealed a slight loss of ST2 resolution within the lung $CD45^+$ population. ILC2s are known to downregulate ST2 expression under specific inflammatory conditions, adopting a $ST2^-KLRG1^{hi}$ phenotype[5], and this loss of resolution will therefore hinder the accurate identification of this subset.

While investigating the spillover between R718 and APC-Cy7, visual inspection revealed significant additional spillover of APC-Cy7 into PE-Cy7 despite the relatively mid-range spread value of 4.87 (Online Table 4), resulting in near complete loss of the TRAIL signal (Online Figure 14A). This spread was also observed to impact PE-Cy5 and BV786 resolution. Such dramatic spread was most likely due to the high antigen density of CD45 coupled with the increased sensitivity of the Cytek® Aurora. However, reducing the concentration of CD45 APC-Cy7 from 1:200 to 1:800 (Online Figure 14B) had little impact on overall spread (data not shown). To account for this, CD45 APC-Cy7 was replaced with CD45 BUV395 due to its significantly reduced brightness, thus improving TRAIL resolution (Online Figure 14C). This CD45 iteration subsequently resulted in a sequential shift in the fluorochromes assigned to both GATA-3 and IL-22. In this regard, GATA-3 was moved from BUV395 to BB700, a much brighter fluorochrome, thus providing enhanced resolution of GATA-$3^+$ cells within both the lung and siLP (Online Figure 15). BB700 and PerCP-Cy5.5 (IL-22) both peak within the B9 emission channel, and hence cannot be used in combination. However, the initial SSM identified a potential issue with PE-Cy7 spreading into PerCP-Cy5.5, and therefore shifting IL-22 to Alexa Fluor 647 allowed the avoidance of this issue, whilst permitting the use of GATA-3 BB700 within the panel (see "Reference control optimisation" for further justification of this iteration).

Lastly, it is important to note this panel has not incorporated the use of BUV536, BUV615, BUV661 and BUV805 on the 355nm ultraviolet laser. These fluorochromes all have tight emissions bands and limited spread, and therefore addition of these fluorochromes allows user-specific expansion of the panel whilst limiting panel Complexity™ to 10.82.

***Reference control optimisation***
Cell and bead comparisons (Online Figure 16) were performed for all reference controls, and compared to the expected spectral profile provided by the Cytek Full Spectrum Viewer (https://spectrum.cytekbio.com/). Cellular fixation is known to alter both the spectral profile of fluorochromes and the autofluorescent properties of cells[6]. To account for this, all reference controls (including unstained reference cells) underwent the exact staining protocol as fully stained samples with regards to fixation/permeabilisation and buffers used, with the exception of BSB+ as this is known to alter the spectral profile of some beads. Cell reference controls were used when a sufficient number of positively labelled cells could be obtained for accurate unmixing. Where it was difficult to obtain a sufficient number of positively stained cells, beads were used for future unmixing, providing the signature was equivalent to the cellular signature or matched the Cytek reference spectra. Consequently, IL-22 needed to be moved off the PerCP-Cy5.5 fluorochrome (Online Figure 17) as sufficient

positive cells could not reliably be collected, the host species is goat and thus will not bind to UltraComp eBeads™, and PerCP-Cy5.5 is a tandem fluorochrome, so a surrogate marker was not able to be used. Moreover, PerCP-Cy5.5 was previously identified as having a high spectral similarity with PerCP-eFluor 710 and PE-Cy5. To account for these issues, IL-22 conjugated to Alexa Fluor 647 was used in the final OMIP and GATA-3 was used as a surrogate antibody for defining an accurate bead reference control.

***Final in silico optimisation***

*In silico* optimisation of the final OMIP iteration containing CD45 BUV395, GATA-3 BB700, LIVE/DEAD™ Blue and IL-22 Alexa Fluor 647 was performed using the SIM (Online Table 5). The SIM identified the highest similarities, and therefore increased risk of spreading, to occur between PerCP-eFluor 710 and BB700 (SI = 0.82) and Alexa Fluor 647 and APC (SI = 0.9; Online Table 5). However, due to careful fluorochrome assignment, the final OMIP avoids co-expressing markers on these fluorochrome pairs, thus minimising this potential issue (also identified by the final SSM; Online Table 6). This consideration is particularly important for the combination of Alexa Fluor 647 and APC (Online Figure 18), which would not be achievable on a conventional cytometer.

***Data analysis***

All manual analysis was performed using FlowJo™ version 10.8.1. Given the breadth and complexity of modern full spectrum (and conventional) panels, computational analysis is routinely used as a complementary method of subset characterisation. These unbiased analysis pipelines additionally allow the discovery of novel subsets which may go unnoticed with manual gating.

To perform high-dimensional unbiased analysis of the panel presented here, all samples were pre-gated as per Figure 1A. Total IL-7R$\alpha^+$Lin$^-$ ILCs were then selected (Figure 1B) and CD49b$^+$ cells removed. Final gates were exported as separate FCS files. Unsupervised analysis was performed in the R environment for statistical computing (version 4.1.2). Following file importation, fluorescent parameters were filtered to remove LIVE/DEAD™ Blue, CD45 BUV395, IL-7R$\alpha$ PE-Cy7, Lin FITC and CD49b eFluor 506 prior to analysis. Data underwent logicle transformation followed by quality assessment with *flowCut*[7] to check for aberrant fluorescent signatures, with all files passing. Dimensionality reduction was performed using the *CATALYST* package[8], employing initial high-resolution *FlowSOM* clustering prior to low-resolution metacluster generation via *ConsensusClusterPlus*. Uniform Manifold Approximation and Projection (UMAP)[9] was performed on 100 cells per sample. ILC-specific cytokines and Ki-67 were excluded from the criteria used for subset clustering. Heatmaps displaying marker expression profiles within each defined cluster were used to immunophenotypically characterise individual clusters (Online Figure 19), prior to subsetting based on tissue localisation (Online Figure 20). Cytokine expression profiles and proliferation was then overlayed on each cluster (Online Figure 21).

**Tissue Preparation**

Female BALB/c mice were autopsied at 8-12 weeks of age and whole lung was collected into individual Bijou containers with RPMI 1640 + 10% fetal calf serum (FCS; R10F; Online Table 3) and kept on ice. Small intestine (SI) was removed by cutting between the stomach and cecum. SI was flushed with Hank's Balanced Salt Solution (HBSS) + 2% FCS, followed by removal of Peyer's Patches and adipose tissue. Cleaned SI was cut longitudinally, flushed of remaining contents with HBSS + 2% FCS, transferred to a Bijou container with R10F and kept on ice.

*Lung single-cell preparation*

1. Aspirate R10F from Bijou collection tube to remove contaminating blood.
2. Manually disaggregate samples with surgical scissors and resuspend in 1ml R10F for enzymatic digestion (0.15mg/ml Liberase and 25μg/ml DNase).
3. Incubate Bijou container at 37°C for 30 minutes under agitation (250rpm).
4. Add 1ml R10F + 5mM EDTA to inactivate the Liberase and leave on ice for 5 minutes.
5. Manually disrupt sample via pipette to dislodge cells from digested tissue, pass through a 70μm filter and wash with 4ml R10F.
6. Centrifuge at 800 x *g* for 5 minutes.
7. Remove supernatant and resuspend pellet in 2ml red blood cell (RBC) lysis buffer (sterile $dH_20$, 0.1% $KHCO_3$, 0.83% $NH_4Cl$, 0.037% EDTA) for 2 minutes.
8. Wash cells with 4ml R10F to inactivate RBC lysis buffer and centrifuge at 800 x *g* for 5 minutes.
9. Remove supernatant and resuspend pellet in 1ml R10F for cell counting.

*Small intestine lamina propria (siLP) single-cell preparation*

1. Cut SI into 1cm pieces.
2. Transfer to 50ml Falcon tube and vortex in 15ml HBSS for 10 seconds.
3. Filter through nylon mesh, discard the HBSS supernatant, transfer SI back to 50ml Falcon tube and add 15ml fresh HBSS and vortex for 10 seconds.
4. Filter through nylon mesh and discard HBSS supernatant.
5. Transfer SI to fresh 50ml Falcon tube and add 15ml epithelial stripping buffer (2mM EDTA, 10mM HEPES, 15ml HBSS).
6. Secure Falcon tubes horizontally in a water bath and incubate at 37°C for 15 minutes under agitation (250rpm).
7. Vortex samples (full speed) for 10 seconds and filter through nylon mesh.
8. Discard supernatant and repeat steps 5-7.
9. Wash SI with 25ml HBSS and vortex (full speed) for 10 seconds to remove residual epithelial stripping buffer.
10. Filter through nylon mesh, discard supernatant and repeat step 9.
11. Transfer SI to fresh 50ml Falcon tube and add 10ml digest buffer (10ml R10F, 1mg/ml Collagenase VIII from *Clostridium histolyticum*, 20μg/ml DNase I).
12. Incubate at 37°C for 30 minutes under agitation (250rpm).
13. Vortex samples (full speed) for 10 seconds and filter through 70μm cell strainer. Wash cell strainer with 2x 5ml R10F.
10. Centrifuge at 800 x *g* for 5 minutes.
11. Remove supernatant and resuspend pellet in 1ml R10F for cell counting.

***Ex vivo* stimulation**

1. Prepare all *ex vivo* stimulation conditions in sterile R10F;
   a. R10F only (unstimulated control)
   b. 5µg/ml Brefeldin A (BFA only)
   c. 20ng/ml PMA + 2µM ionomycin + 5µg/ml Brefeldin A (base stimulation cocktail)
   d. pro-ILC1 = base stimulation cocktail + rmIL-12 (20ng/ml) + rmIL-18 (20ng/ml)
   e. pro-ILC2 = base stimulation cocktail + rmIL-33 (20ng/ml)
   f. pro-ILC3 = base stimulation cocktail + rmIL-1β (20ng/ml) + rmIL-23 (20ng/ml)
2. Add $1x10^6$ single cell suspension for each sample to a 96 well round bottom culture plate.
3. Include an additional well of $1x10^6$ single cell suspension for each tissue type for the unstained control and additional wells (excess lung routinely used) for the fluorochrome reference controls.
4. Add 1 drop of UltraComp eBeads™ Compensation Beads to required wells for fluorochrome reference controls.
5. Centrifuge at 800 x *g* for 5 minutes at 4°C.
6. Remove supernatant and add 200µl of *ex vivo* stimulation conditions (per 1a-f) to appropriate wells.
7. Add 200µl base stimulation cocktail to UltraComp eBeads™ wells (NB. Fluorochrome reference controls must be treated exactly the same as samples).
8. Incubate at 37°C in 5% $CO_2$ for 5 hours.
9. Centrifuge at 800 x *g* for 5 minutes at 4°C.
10. Remove supernatant and wash pellet with 150µl PBS, followed by centrifugation at 800 x *g* for 5 minutes at 4°C.
11. Repeat step 10.

**Staining Protocol**

1. Resuspend cells in 100µl Viability LIVE/DEAD™ Fixable Blue. Unstained controls and reference controls are resuspended in 100µl Flow Buffer (PBS + 0.1% BSA + 10% sodium azide). Incubate for 15 minutes at room temperature in the dark.
2. Wash wells with 150µl Flow Buffer, followed by centrifugation at 800 x *g* for 5 minutes at 4°C.
3. Remove supernatant and repeat step 2.
4. Remove supernatant and resuspend cells in 30µl Fc Block™. Resuspend UltraComp eBeads™ in 30µl Flow Buffer. Incubate for 15 minutes at 4°C in the dark.
5. Add 30µl extracellular staining cocktail + Brilliant Stain Buffer Plus to samples or respective fluorescently conjugated antibody to single stain cell controls, for a final volume of 60µl/$1x10^6$ cells. Add appropriate fluorochrome to UltraComp eBeads™. Stain for 30 minutes at 4°C in the dark.
6. Wash wells with 150µl Flow Buffer, followed by centrifugation at 800 x *g* for 5 minutes at 4°C.
7. Remove supernatant and repeat step 6.
8. Remove supernatant and resuspend all wells in 100µl fixation/permeabilization buffer (Foxp3/Transcription Factor Staining Buffer Set) for 30 minutes at 4°C in the dark.
9. Wash wells with 100µl 1x permeabilization buffer, followed by centrifugation at 800 x *g* for 5 minutes at 4°C.
10. Remove supernatant and repeat step 9.

11. Remove supernatant and resuspend samples in 30µl intracellular staining cocktail. Resuspend unstained controls and reference controls in 1x permeabilization buffer. Incubate for 60 minutes at 4°C in the dark.
12. Wash wells with 150µl 1x permeabilization buffer, followed by centrifugation at 800 x *g* for 5 minutes at 4°C.
13. Remove supernatant and repeat step 12.
14. Remove supernatant and resuspend wells in 200µl Flow Buffer
15. Acquire samples on a 5-laser Cytek® Aurora.

**Online Table 1. Instrument optical configuration.** The 5 laser Cytek® Aurora is equipped with a total of 16 detectors for the 355nm ultraviolet (UV) laser, 16 detectors for the 405nm violet laser, 14 detectors for the 488nm blue laser, 10 detectors for the 561nm yellow-green laser and 8 detectors for the 640nm red laser. All fluorochromes listed are those used in the final OMIP.

| Laser | Power | Channel | Centre Wavelength (nm) | Bandwidth (nm) | Parameter |
|---|---|---|---|---|---|
| Ultra Violet (355nm) | 29mW | UV2 | 387 | 15 | BUV395 |
| | | UV6 | 473 | 15 | LIVE/DEAD™ Blue |
| | | UV7 | 514 | 28 | BUV496 |
| | | UV14 | 750 | 30 | BUV737 |
| Violet (405nm) | 100mW | V1 | 428 | 15 | BV421 |
| | | V3 | 458 | 15 | eFluor 450 |
| | | V5 | 508 | 20 | BV480 |
| | | V7 | 542 | 17 | eFluor 506 |
| | | V11 | 664 | 27 | BV650 |
| | | V13 | 730 | 29 | BV711 |
| | | V15 | 780 | 30 | BV786 |
| Blue (488nm) | 50mW | B2 | 525 | 17 | FITC |
| | | B3 | 542 | 17 | Alexa Fluor 532 |
| | | B9 | 697 | 19 | BB700 |
| | | B10 | 717 | 20 | PerCP-eFluor 710 |
| Yellow-Green (561nm) | 50mW | YG1 | 577 | 20 | PE |
| | | YG3 | 615 | 20 | PE/Dazzle 594 |
| | | YG5 | 678 | 18 | PE-Cy5 |
| | | YG9 | 780 | 30 | PE-Cy7 |
| Red (640nm) | 80mW | R1 | 660 | 17 | APC |
| | | R4 | 717 | 20 | Alexa Fluor 647 |
| | | R7 | 783 | 23 | R718 |

**Online Table 2. Reagents used in final OMIP**

| Specificity | Fluorochrome | Clone | Catalog number | Vendor | Titration |
|---|---|---|---|---|---|
| CD45 | BUV395 | 30-F11 | 565967 | BD Biosciences | 1:200 |
| Dead cells | LIVE/DEAD™ Blue | - | L34961 | eBioscience | 1:2000 |
| CD49a | BUV496 | Ha31/8 | 741111 | BD Biosciences | 1:200 |
| IFN-γ | BUV737 | XMG1.2 | 612769 | BD Biosciences | 1:80 |
| T-bet | BV421 | 4B10 | 644816 | BioLegend | 1:40 |
| IL-13 | eFluor 450 | eBio13A | 48-7133-82 | eBioscience | 1:80 |
| KLRG1 | BV480 | 2F1 | 746353 | BD Biosciences | 1:100 |
| CD49b | eFluor 506 | DX5 | 69-5971-82 | eBioscience | 1:100 |
| IL-17A | BV650 | TC11-18H10.1 | 506929 | BD Biosciences | 1:80 |
| CCR6 | BV711 | 140706 | 740646 | BD Biosciences | 1:200 |
| RORγt | BV786 | Q31-378 | 564723 | BD Biosciences | 1:80 |
| B220 | FITC | RA3-6B2 | 103206 | BioLegend | 1:100 |
| CD3 | FITC | 145-2C11 | 100306 | BioLegend | 1:400 |
| CD4 | FITC | GK1.5 | 100406 | BioLegend | 1:200 |
| CD5 | FITC | 53-7.3 | 100606 | BioLegend | 1:800 |
| CD11b | FITC | M1/70 | 101206 | BioLegend | 1:400 |
| CD11c | FITC | N418 | 117306 | BioLegend | 1:400 |
| CD19 | FITC | 1D3/CD19 | 152404 | BioLegend | 1:400 |
| F4/80 | FITC | BM8 | 123108 | BioLegend | 1:100 |
| FcεR1 | FITC | MAR-1 | 134306 | BioLegend | 1:100 |
| Gr-1 | FITC | RB6-85C | 108406 | BioLegend | 1:200 |
| TCRβ | FITC | H57-597 | 109206 | BioLegend | 1:800 |
| TCRγδ | FITC | UC7-13D5 | 107504 | BioLegend | 1:400 |
| Ter119 | FITC | TER-119 | 116206 | BioLegend | 1:200 |
| Ki-67 | Alexa Fluor 532 | SolA15 | 58-5698-82 | eBioscience | 1:320 |
| GATA-3 | BB700 | L50-823 | 566642 | BD Biosciences | 1:160 |
| IL-18Rα | PerCP-eFluor 710 | P3TUNYA | 46-5183-82 | eBioscience | 1:200 |
| PLZF | PE | MAGS21F7 | 12-9320-80 | eBioscience | 1:320 |
| NKp46 | PE/Dazzle 594 | 29A1.4 | 137630 | BioLegend | 1:100 |
| IL-7Rα | PE-Cy5 | SB/199 | 135014 | BioLegend | 1:200 |
| TRAIL | PE-Cy7 | N2B2 | 109312 | BioLegend | 1:200 |
| IL-5 | APC | TRFK5 | 504306 | BioLegend | 1:20 |
| IL-22 | Alexa Fluor 647 | Poly5164 | 516406 | BioLegend | 1:20 |
| ST2 | R718 | U29-93 | 752185 | BD Biosciences | 1:100 |

**Online Table 3. Reagents used in panel design and optimisation**

| Reagent | Cat. Number | Vendor |
|---|---|---|
| Fetal calf serum | 16000044 | Gibco |
| Bovine serum albumin | A9418-500G | Merck |
| Phosphate buffered saline (PBS) | 437117K | VWR Chemical |
| Liberase | 5401127001 | Roche |
| Collagenase VIII | C2139 | Roche |
| DNase I | 10104159001 | Roche |
| Ammonium chloride ($NH_4Cl$) | 21236.267 | VWR Chemical |
| Potassium bicarbonate ($KHCO_3$) | 237205 | Merck |
| EDTA | 280214S | VWR Chemical |
| RPMI-1640 + L-Glutamine | 21875091 | ThermoFisher Scientific |
| Hank's Balanced Salt Solution (HBSS) | 14185045 | ThermoFisher Scientific |
| Phorbol 12-myrisate 13-acetate (PMA) | P1585-1MG | Sigma-Aldrich |
| Ionomycin | 407950-5 | Merck Millipore |
| Brefeldin A | BML-G405-0005 | Enzo Life Sciences |
| Recombinant Mouse IL-12 (p70) (carrier free) | 577006 | BioLegend |
| Recombinant Mouse IL-18 (carrier free) | 767006 | BioLegend |
| Recombinant Mouse IL-1b (carrier free) | 575106 | BioLegend |
| Recombinant Mouse IL-23 (carrier free) | 589006 | BioLegend |
| Recombinant Mouse IL-33 (carrier free) | 580506 | BioLegend |
| UltraPure™ 0.5M EDTA | 15575020 | ThermoFisher Scientific |
| HEPES | 15630080 | ThermoFisher Scientific |
| Brilliant Stain Buffer Plus (BSB+) | 566385 | BD Biosciences |
| Foxp3/Transcription Factor Staining Buffer Set | 00-5523-00 | eBioscience |
| SpectroFlo® QC Beads | N7-97355 | Cytek Biosciences |
| UltraComp eBeads™ Compensation Beads | 01-2222-42 | ThermoFisher Scientific |
| GATA-3 BUV395 | 565448 | BD Biosciences |
| IL-22 PerCP-Cy5.5 | 516411 | BioLegend |
| GATA-3 Alexa Fluor 647 | 560068 | BD Bioscience |
| CD45 APC-Cy7 | 103116 | BioLegend |
| Fc Block™ | 553142 | BD Biosciences |

**Online Table 4. Spillover Spread Matrix (SSM) of original panel containing CD45 APC-Cy7, GATA-3 BUV395, IL-22 PerCP-Cy5.5 and FVS 575V.**

| | APC | APC-Cy7 | Alexa Fluor 532 | BUV395 | BUV496 | BUV737 | BV421 | BV480 | BV650 | BV711 | BV786 | FITC | FVS 575V | PE | PE-Cy5 | PE-Cy7 | PE/Dazzle 594 | PerCP-Cy5.5 | PerCP-eFluor 710 | R718 | eFluor 450 | eFluor 506 |
|---|---|---|---|---|---|---|---|---|---|---|---|---|---|---|---|---|---|---|---|---|---|---|
| APC | | 1.28 | 0.00 | 0.00 | 0.00 | 0.00 | 0.00 | 0.00 | 1.90 | 0.00 | 0.00 | 0.00 | 0.00 | 0.00 | 0.00 | 0.00 | 0.00 | 1.62 | 0.69 | 1.19 | 0.00 | 0.00 |
| APC-Cy7 | 3.03 | | 1.36 | 0.83 | 2.50 | 2.37 | 0.62 | 2.35 | 1.21 | 2.10 | 1.81 | 0.45 | 1.88 | 0.52 | 1.50 | 4.87 | 1.82 | 1.92 | 1.72 | 2.51 | 1.66 | 1.60 |
| Alexa Fluor 532 | 0.39 | 0.00 | | 0.00 | 1.06 | 0.42 | 0.00 | 0.00 | 1.36 | 0.50 | 0.40 | 0.94 | 4.27 | 4.72 | 1.51 | 0.46 | 3.19 | 0.66 | 0.94 | 0.66 | 1.63 | 2.89 |
| BUV395 | 0.00 | 0.00 | 0.47 | | 2.96 | 0.35 | 0.30 | 1.36 | 0.47 | 0.00 | 0.40 | 1.23 | 0.73 | 0.64 | 0.78 | 0.00 | 0.84 | 0.39 | 0.00 | 0.40 | 0.51 | 1.02 |
| BUV496 | 0.69 | 0.54 | 2.12 | 1.61 | | 0.85 | 0.88 | 5.39 | 1.86 | 1.10 | 0.85 | 2.21 | 3.61 | 1.11 | 1.58 | 0.53 | 3.50 | 1.24 | 1.38 | 0.78 | 2.70 | 4.36 |
| BUV737 | 0.14 | 1.78 | 0.75 | 0.44 | 2.25 | | 0.18 | 0.00 | 0.77 | 1.26 | 1.39 | 0.00 | 2.08 | 0.31 | 0.32 | 0.70 | 1.42 | 2.29 | 1.67 | 2.68 | 0.49 | 1.38 |
| BV421 | 1.60 | 0.75 | 3.18 | 2.22 | 4.80 | 0.92 | | 8.40 | 1.94 | 1.68 | 1.19 | 3.31 | 4.66 | 2.13 | 1.90 | 0.52 | 3.17 | 1.86 | 1.66 | 0.79 | 5.24 | 5.94 |
| BV480 | 0.17 | 0.00 | 1.01 | 0.29 | 4.14 | 0.35 | 0.34 | | 1.15 | 0.35 | 0.36 | 1.03 | 2.41 | 0.38 | 0.90 | 0.18 | 1.51 | 0.26 | 0.35 | 0.30 | 0.55 | 2.82 |
| BV650 | 0.89 | 0.51 | 0.86 | 0.19 | 2.41 | 1.02 | 1.06 | 2.78 | | 2.20 | 1.19 | 1.12 | 2.24 | 0.46 | 4.58 | 0.49 | 1.72 | 2.60 | 0.99 | 1.23 | 0.92 | 1.86 |
| BV711 | 0.40 | 1.54 | 1.27 | 0.95 | 2.18 | 2.35 | 1.29 | 4.17 | 1.40 | | 2.63 | 1.15 | 3.51 | 0.73 | 1.13 | 0.72 | 2.59 | 2.40 | 2.07 | 2.29 | 2.39 | 2.43 |
| BV786 | 0.18 | 1.34 | 0.59 | 0.21 | 1.44 | 1.69 | 2.85 | 1.28 | 0.62 | 2.11 | | 12.55 | 1.17 | 0.00 | 0.82 | 0.99 | 0.85 | 0.50 | 0.36 | 0.72 | 0.75 | 1.15 |
| FITC | 0.66 | 0.73 | 5.82 | 2.01 | 4.89 | 1.24 | 1.30 | 7.64 | 2.12 | 1.38 | 1.28 | | 5.79 | 1.93 | 2.97 | 0.50 | 3.31 | 1.44 | 1.22 | 0.93 | 3.96 | 4.14 |
| FVS 575V | 0.53 | 0.29 | 1.62 | 1.07 | 3.42 | 0.83 | 0.82 | 4.75 | 2.68 | 1.48 | 1.28 | 1.99 | | 0.91 | 1.71 | 0.50 | 1.89 | 0.92 | 0.69 | 0.75 | 1.73 | 2.83 |
| PE | 0.00 | 0.00 | 0.03 | 0.00 | 0.55 | 0.00 | 0.00 | 0.00 | 2.26 | 0.52 | 0.70 | 0.00 | 6.02 | | 2.14 | 0.62 | 4.00 | 0.00 | 1.01 | 0.74 | 0.00 | 3.52 |
| PE-Cy5 | 1.66 | 1.20 | 3.92 | 0.89 | 1.93 | 0.80 | 0.51 | 6.59 | 4.50 | 2.55 | 1.34 | 2.53 | 10.48 | 1.46 | | 2.30 | 6.03 | 7.77 | 3.59 | 2.75 | 2.53 | 4.30 |
| PE-Cy7 | 0.61 | 1.81 | 1.72 | 1.23 | 4.11 | 1.49 | 0.94 | 5.99 | 2.24 | 3.01 | 1.87 | 1.64 | 5.83 | 1.51 | 3.56 | | 4.59 | 1.02 | 1.91 | 0.96 | 2.91 | 3.23 |
| PE/Dazzle 594 | 0.58 | 0.45 | 0.04 | 0.73 | 0.00 | 0.00 | 0.72 | 0.00 | 2.45 | 1.09 | 0.71 | 2.47 | 7.47 | 2.76 | 3.48 | 1.11 | | 1.81 | 1.76 | 0.91 | 2.00 | 4.17 |
| PerCP-Cy5.5 | 1.47 | 2.88 | 0.07 | 1.71 | 5.02 | 0.00 | 0.00 | 0.00 | 0.00 | 3.37 | 0.00 | 0.00 | 0.00 | 3.18 | 4.37 | 2.26 | 3.74 | | 1.95 | 2.20 | 0.00 | 6.93 |
| PerCP-eFluor 710 | 0.80 | 3.37 | 2.71 | 0.16 | 2.62 | 2.65 | 0.86 | 2.58 | 2.79 | 4.46 | 2.22 | 1.09 | 8.65 | 1.20 | 2.76 | 3.21 | 5.46 | 9.62 | | 10.83 | 2.37 | 6.60 |
| R718 | 0.42 | 10.44 | 0.56 | 0.20 | 1.58 | 1.98 | 0.26 | 0.00 | 0.55 | 1.68 | 1.09 | 0.39 | 1.76 | 0.34 | 0.53 | 2.36 | 0.99 | 4.21 | 2.46 | | 0.34 | 0.77 |
| eFluor 450 | 0.00 | 0.22 | 0.42 | 0.00 | 1.69 | 0.00 | 1.02 | 4.70 | 1.19 | 0.00 | 0.24 | 0.71 | 2.06 | 0.25 | 0.68 | 0.21 | 1.53 | 0.29 | 0.45 | 0.39 | | 3.72 |
| eFluor 506 | 0.00 | 0.32 | 1.51 | 0.01 | 1.60 | 0.45 | 0.37 | 3.53 | 0.93 | 0.00 | 0.72 | 1.34 | 2.41 | 0.81 | 0.76 | 0.00 | 1.30 | 0.00 | 0.37 | 0.38 | 0.62 | |

**Online Table 5. Similarity Index Matrix (SIM) of final OMIP.**

| | BUV395 | LIVE/DEAD Blue | BUV496 | BUV737 | BV421 | eFluor 450 | BV480 | eFluor 506 | BV650 | BV711 | BV786 | FITC | Alexa Fluor 532 | BB700 | PerCP-eFluor 710 | PE | PE/Dazzle 594 | PE-Cy5 | PE-Cy7 | APC | Alexa Fluor 647 | R718 |
|---|---|---|---|---|---|---|---|---|---|---|---|---|---|---|---|---|---|---|---|---|---|---|
| BUV395 | 1 | | | | | | | | | | | | | | | | | | | | | |
| LIVE/DEAD Blue | 0.51 | 1 | | | | | | | | | | | | | | | | | | | | |
| BUV496 | 0.25 | 0.58 | 1 | | | | | | | | | | | | | | | | | | | |
| BUV737 | 0.03 | 0.02 | 0.01 | 1 | | | | | | | | | | | | | | | | | | |
| BV421 | 0.05 | 0.26 | 0.09 | 0 | 1 | | | | | | | | | | | | | | | | | |
| eFluor 450 | 0 | 0.21 | 0.08 | 0 | 0.85 | 1 | | | | | | | | | | | | | | | | |
| BV480 | 0.02 | 0.2 | 0.38 | 0 | 0.28 | 0.46 | 1 | | | | | | | | | | | | | | | |
| eFluor 506 | 0.01 | 0.07 | 0.27 | 0 | 0.14 | 0.24 | 0.87 | 1 | | | | | | | | | | | | | | |
| BV650 | 0 | 0.02 | 0.02 | 0.13 | 0.09 | 0.08 | 0.06 | 0.05 | 1 | | | | | | | | | | | | | |
| BV711 | 0 | 0.02 | 0 | 0.41 | 0.09 | 0.08 | 0.03 | 0.02 | 0.45 | 1 | | | | | | | | | | | | |
| BV786 | 0 | 0.04 | 0.01 | 0.29 | 0.2 | 0.17 | 0.05 | 0.03 | 0.19 | 0.54 | 1 | | | | | | | | | | | |
| FITC | 0.01 | 0.01 | 0.09 | 0 | 0.01 | 0.01 | 0.1 | 0.14 | 0 | 0 | 0 | 1 | | | | | | | | | | |
| Alexa Fluor 532 | 0.03 | 0.02 | 0.02 | 0 | 0.02 | 0.02 | 0.01 | 0.01 | 0.01 | 0 | 0 | 0.55 | 1 | | | | | | | | | |
| BB700 | 0 | 0 | 0 | 0.3 | 0 | 0 | 0.02 | 0.02 | 0.37 | 0.53 | 0.22 | 0.02 | 0.07 | 1 | | | | | | | | |
| PerCP-eFluor 710 | 0 | 0 | 0 | 0.39 | 0 | 0 | 0 | 0 | 0.31 | 0.65 | 0.28 | 0 | 0.02 | 0.82 | 1 | | | | | | | |
| PE | 0.01 | 0 | 0.03 | 0 | 0 | 0.01 | 0.05 | 0.07 | 0.05 | 0.01 | 0 | 0.09 | 0.4 | 0.06 | 0.03 | 1 | | | | | | |
| PE/Dazzle 594 | 0.01 | 0 | 0.02 | 0.02 | 0 | 0 | 0.04 | 0.04 | 0.16 | 0.04 | 0.01 | 0.06 | 0.3 | 0.18 | 0.16 | 0.67 | 1 | | | | | |
| PE-Cy5 | 0 | 0 | 0 | 0.1 | 0 | 0 | 0 | 0 | 0.29 | 0.14 | 0.03 | 0.01 | 0.06 | 0.48 | 0.47 | 0.14 | 0.43 | 1 | | | | |
| PE-Cy7 | 0 | 0 | 0 | 0.14 | 0 | 0 | 0 | 0 | 0.04 | 0.11 | 0.17 | 0 | 0.01 | 0.17 | 0.29 | 0.02 | 0.05 | 0.14 | 1 | | | |
| APC | 0 | 0.01 | 0 | 0.24 | 0 | 0 | 0 | 0 | 0.37 | 0.23 | 0.04 | 0 | 0 | 0.33 | 0.29 | 0.04 | 0.16 | 0.52 | 0.05 | 1 | | |
| Alexa Fluor 647 | 0 | 0.01 | 0 | 0.22 | 0 | 0 | 0 | 0 | 0.18 | 0.18 | 0.01 | 0 | 0 | 0.33 | 0.23 | 0.01 | 0.06 | 0.32 | 0.03 | 0.9 | 1 | |
| R718 | 0 | 0 | 0 | 0.52 | 0 | 0 | 0 | 0 | 0.12 | 0.35 | 0.09 | 0.01 | 0.01 | 0.34 | 0.39 | 0 | 0.02 | 0.16 | 0.08 | 0.47 | 0.52 | 1 |

Complexity Index: 10.06

**Online Table 6. Spillover Spread Matrix (SSM) of final OMIP.**

| | APC | Alexa Fluor 532 | Alexa Fluor 647 | BB700 | BUV395 | BUV496 | BUV737 | BV421 | BV480 | BV650 | BV711 | BV786 | FITC | LIVE/DEAD Blue | PE | PE-Cy5 | PE-Cy7 | PE/Dazzle 594 | PerCP-eFluor 710 | R718 | eFluor 450 | eFluor 506 |
|---|---|---|---|---|---|---|---|---|---|---|---|---|---|---|---|---|---|---|---|---|---|---|
| APC | | 0.00 | 4.71 | 0.91 | 0.00 | 0.00 | 0.00 | 0.00 | 0.00 | 1.63 | 0.00 | 0.00 | 0.00 | 0.00 | 0.00 | 0.65 | 0.00 | 0.00 | 1.06 | 1.46 | 0.00 | 0.00 |
| Alexa Fluor 532 | 0.69 | | 1.10 | 1.00 | 0.00 | 0.00 | 0.00 | 0.35 | 0.00 | 0.00 | 0.34 | 0.00 | 1.43 | 0.00 | 2.14 | 0.59 | 0.00 | 0.70 | 0.53 | 0.46 | 0.00 | 0.00 |
| Alexa Fluor 647 | 3.39 | 0.22 | | 0.53 | 0.00 | 2.29 | 0.77 | 0.00 | 0.00 | 0.56 | 0.26 | 0.59 | 0.33 | 0.00 | 0.37 | 1.13 | 0.39 | 0.22 | 0.72 | 1.79 | 0.00 | 0.00 |
| BB700 | 2.82 | 0.94 | 4.02 | | 0.15 | 1.95 | 1.23 | 1.41 | 1.67 | 2.89 | 6.03 | 1.68 | 0.53 | 0.00 | 0.24 | 0.99 | 1.30 | 0.60 | 3.51 | 2.64 | 0.74 | 1.06 |
| BUV395 | 0.00 | 0.00 | 0.00 | 0.00 | | 2.11 | 0.00 | 0.00 | 0.00 | 0.00 | 0.00 | 0.00 | 0.00 | 2.37 | 0.00 | 0.00 | 0.00 | 0.00 | 0.00 | 0.00 | 0.00 | 0.00 |
| BUV496 | 0.97 | 1.63 | 0.97 | 0.90 | 1.22 | | 1.07 | 1.03 | 4.04 | 1.38 | 0.95 | 1.07 | 2.14 | 1.72 | 0.81 | 1.03 | 0.73 | 1.82 | 0.76 | 0.68 | 3.41 | 3.42 |
| BUV737 | 1.16 | 0.17 | 1.52 | 0.83 | 0.72 | 3.50 | | 0.34 | 0.26 | 0.58 | 2.21 | 1.86 | 0.00 | 0.52 | 0.16 | 0.77 | 0.71 | 0.26 | 3.35 | 3.20 | 0.36 | 0.67 |
| BV421 | 0.42 | 0.25 | 0.00 | 0.21 | 0.27 | 1.99 | 0.21 | | 4.66 | 0.81 | 0.00 | 0.21 | 0.58 | 0.87 | 0.00 | 0.28 | 0.12 | 0.40 | 0.28 | 0.00 | 2.88 | 3.02 |
| BV480 | 0.62 | 0.00 | 0.65 | 0.00 | 0.00 | 2.32 | 0.00 | 1.21 | | 1.00 | 0.00 | 0.00 | 0.00 | 1.26 | 0.77 | 0.02 | 0.00 | 1.30 | 0.01 | 0.49 | 0.00 | 4.15 |
| BV650 | 1.76 | 0.30 | 1.53 | 1.09 | 0.14 | 4.26 | 1.15 | 1.57 | 2.25 | | 1.74 | 1.29 | 0.45 | 0.14 | 0.21 | 1.25 | 0.41 | 0.40 | 0.74 | 0.67 | 0.93 | 1.45 |
| BV711 | 1.16 | 1.28 | 1.72 | 1.31 | 1.12 | 3.57 | 2.50 | 1.21 | 3.70 | 1.75 | | 2.95 | 1.79 | 1.57 | 0.89 | 1.26 | 0.80 | 2.16 | 1.47 | 2.37 | 4.13 | 2.75 |
| BV786 | 0.43 | 0.00 | 0.55 | 0.24 | 0.00 | 1.81 | 2.19 | 7.04 | 1.41 | 0.50 | 2.55 | | 0.21 | 0.29 | 0.18 | 0.00 | 0.72 | 0.22 | 0.38 | 1.30 | 0.86 | 0.93 |
| FITC | 0.23 | 2.64 | 0.30 | 0.42 | 0.00 | 1.56 | 0.09 | 0.13 | 1.39 | 0.40 | 0.18 | 0.15 | | 0.00 | 0.75 | 0.45 | 0.07 | 0.39 | 0.25 | 0.15 | 0.16 | 1.45 |
| LIVE/DEAD Blue | 0.59 | 0.00 | 1.49 | 0.97 | 2.88 | 0.00 | 0.00 | 1.37 | 0.00 | 0.64 | 1.02 | 0.00 | 0.00 | | 0.79 | 0.00 | 0.00 | 0.00 | 0.00 | 0.56 | 0.00 | 0.00 |
| PE | 0.68 | 2.35 | 0.31 | 0.45 | 0.00 | 0.00 | 0.00 | 0.00 | 0.00 | 1.15 | 0.00 | 0.00 | 1.06 | 0.00 | | 0.55 | 0.00 | 1.50 | 0.42 | 0.00 | 0.00 | 1.38 |
| PE-Cy5 | 2.95 | 0.36 | 4.62 | 9.62 | 0.18 | 2.24 | 0.78 | 0.51 | 4.47 | 3.05 | 1.47 | 0.48 | 0.88 | 0.11 | 1.07 | | 1.53 | 0.54 | 3.54 | 1.62 | 0.87 | 2.68 |
| PE-Cy7 | 1.28 | 2.84 | 1.58 | 1.47 | 2.17 | 8.01 | 2.23 | 1.58 | 7.54 | 2.53 | 1.42 | 2.17 | 4.54 | 3.50 | 1.62 | 1.82 | | 3.38 | 1.54 | 1.55 | 7.08 | 4.57 |
| PE/Dazzle 594 | 1.84 | 0.61 | 1.71 | 1.93 | 0.00 | 0.00 | 0.00 | 0.57 | 0.00 | 0.88 | 0.40 | 0.00 | 1.09 | 0.00 | 1.59 | 1.68 | 0.55 | | 1.14 | 0.86 | 0.00 | 0.96 |
| PerCP-eFluor 710 | 1.93 | 0.47 | 2.65 | 5.21 | 0.29 | 3.50 | 1.79 | 0.44 | 0.00 | 1.43 | 3.70 | 1.94 | 0.48 | 0.29 | 0.28 | 2.08 | 2.12 | 0.25 | | 7.49 | 0.39 | 0.94 |
| R718 | 4.76 | 0.30 | 9.30 | 2.03 | 0.00 | 3.73 | 2.84 | 0.00 | 0.90 | 1.57 | 1.80 | 1.20 | 0.00 | 0.23 | 0.22 | 0.84 | 0.82 | 0.19 | 3.11 | | 1.16 | 0.72 |
| eFluor 450 | 0.28 | 0.37 | 0.00 | 0.17 | 0.00 | 2.05 | 0.33 | 1.02 | 5.40 | 0.88 | 0.00 | 0.23 | 0.73 | 0.67 | 0.00 | 0.27 | 0.28 | 0.49 | 0.00 | 0.00 | | 3.69 |
| eFluor 506 | 0.88 | 1.26 | 0.87 | 0.53 | 0.00 | 2.72 | 0.00 | 0.55 | 4.24 | 0.46 | 0.00 | 0.00 | 1.95 | 0.00 | 0.51 | 0.00 | 0.25 | 0.63 | 0.01 | 0.59 | 0.72 | |

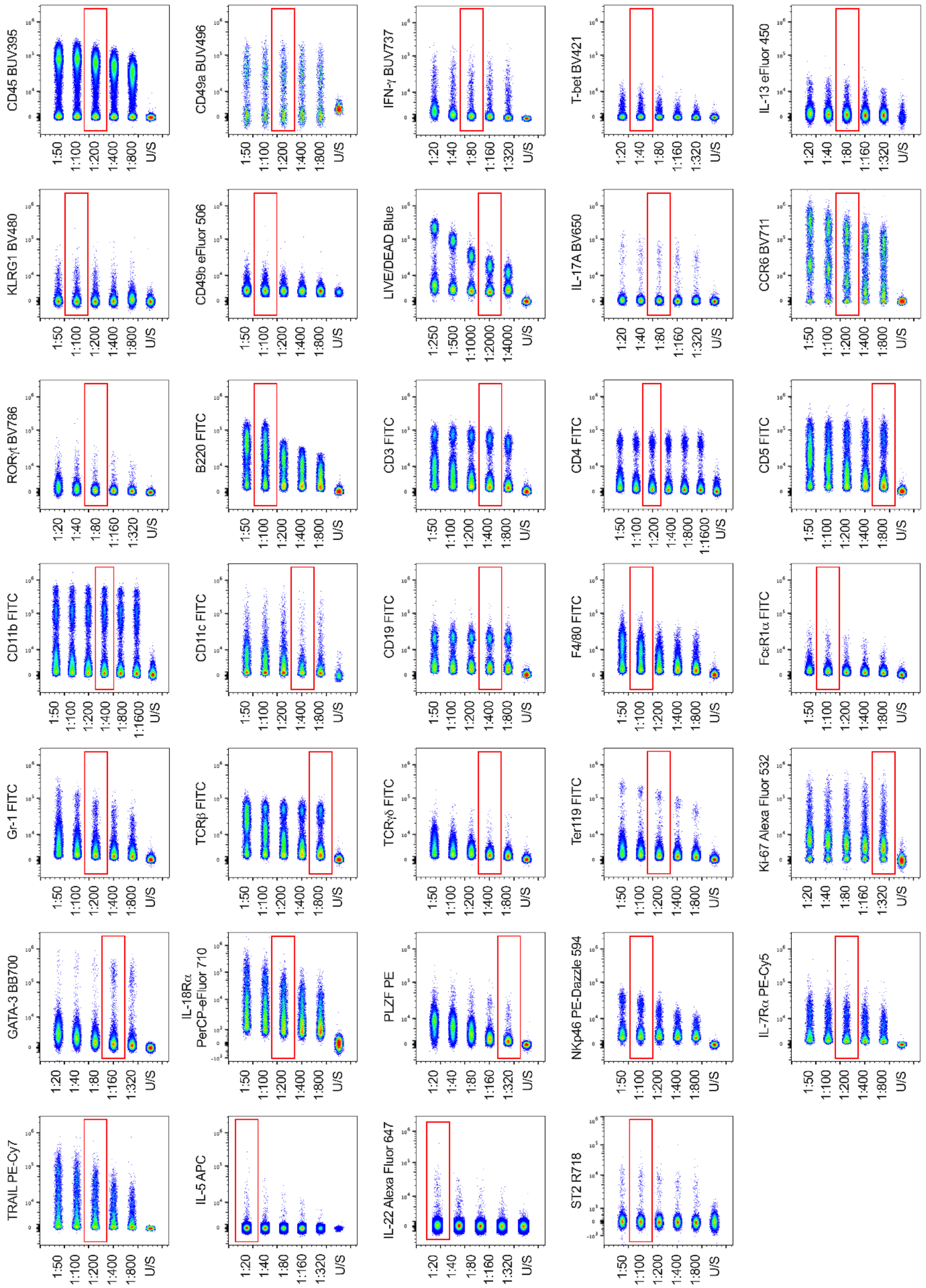


**Online Figure 1. Antibody titrations used in the final OMIP.** All antibodies were individually titrated on lung single cells from BALB/c mice to determine their optimal concentration. All single cell suspensions were stimulated *ex vivo* with PMA, Ionomycin and Brefeldin A (base stimulation cocktail) for 5 hours prior to staining. To obtain a sufficient positive signal for cytokine titrations, appropriate single cells were additionally stimulated with IL-12 + IL-18 (IFN-γ BUV737; pro-ILC1 stimulation cocktail), IL-33 (IL-5 APC and IL-13 eFluor 450; pro-ILC2 stimulation cocktail) or IL-1β + IL-23 (IL-17A BV650 and IL-22 Alexa Fluor 647; pro-ILC3 stimulation cocktail). Data are pre-gated to remove cellular debris (SSC v FSC), RBC contamination (SSC-A v SSC-B-A) and doublets (FSC-A v FSC-H). U/S = unstained control. Red boxes indicate dilution used in the final OMIP.

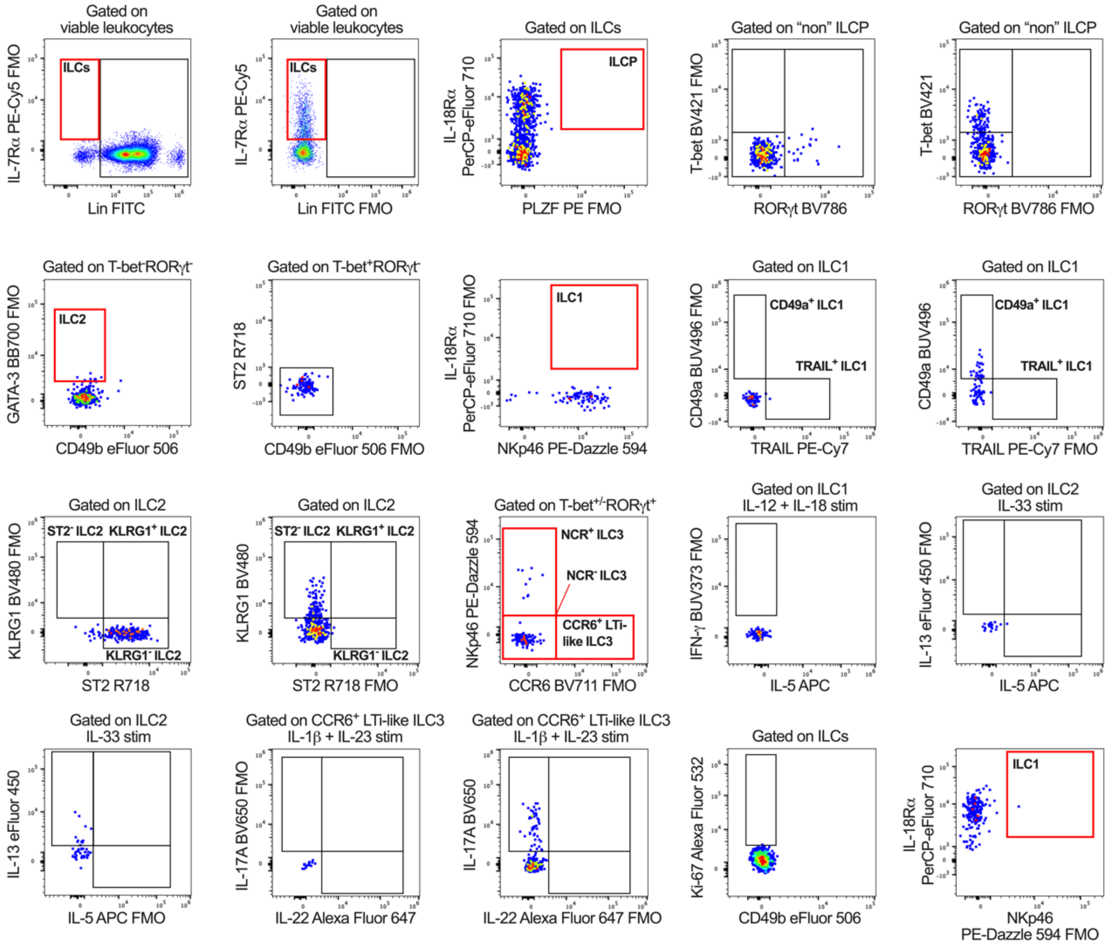


**Online Figure 2. Fluorescence minus one (FMO) controls.** Data are lung single cells showing FMO controls for intermediate (black) and terminal (red) gates displayed in Figure 1 (where required). Single cell suspensions were stimulated *ex vivo* with PMA, Ionomycin and Brefeldin A for 5 hours prior to staining.

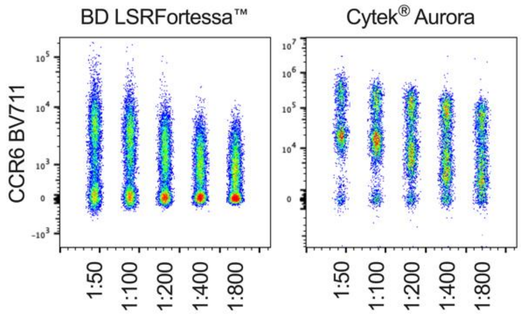

**Online Figure 3. Antibody titration comparison between conventional and full spectrum flow cytometry.** Lung single cells demonstrating fluorescence of CCR6 BV711 titrated on a BD LSRFortessa™ and Cytek® Aurora. Single cell suspensions were stimulated *ex vivo* with PMA, Ionomycin and Brefeldin A (base stimulation cocktail) for 5 hours prior to staining. Data are pre-gated to remove cellular debris (SSC v FSC), RBC contamination (SSC-A v SSC-B-A) and doublets (FSC-A v FSC-H).

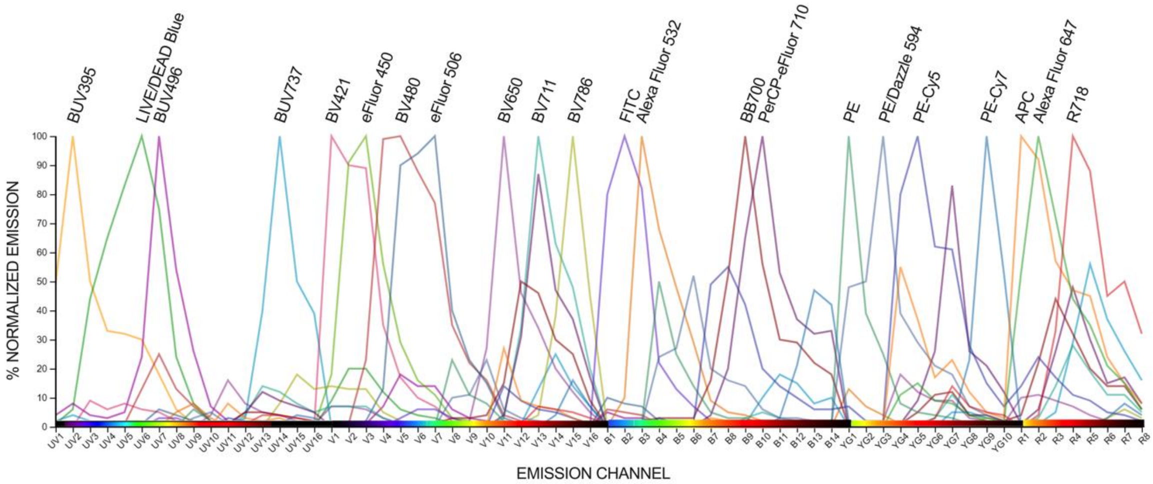


**Online Figure 4. Fluorochrome emission spectra.** Individual emission spectra for all 22 fluorochromes used in the final OMIP.

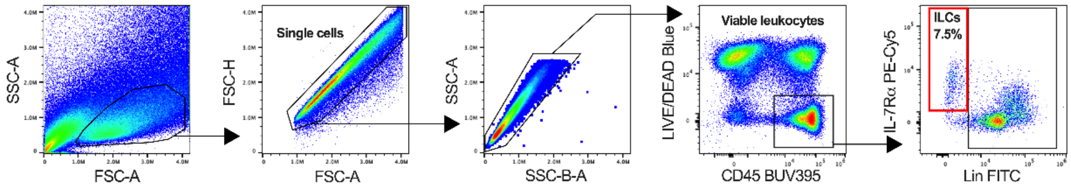


**Online Figure 5. Initial gating strategy to define siLP ILCs.** After exclusion of cellular debris, doublets, red blood cell contamination and non-viable CD45$^{+/-}$ cells, viable CD45$^{+}$ siLP leukocytes were gated on IL-7R$\alpha$ and lineage (Lin) negative cells to identify ILCs. Single cell suspensions were stimulated *ex vivo* with PMA, Ionomycin and Brefeldin A for 5 hours prior to staining.

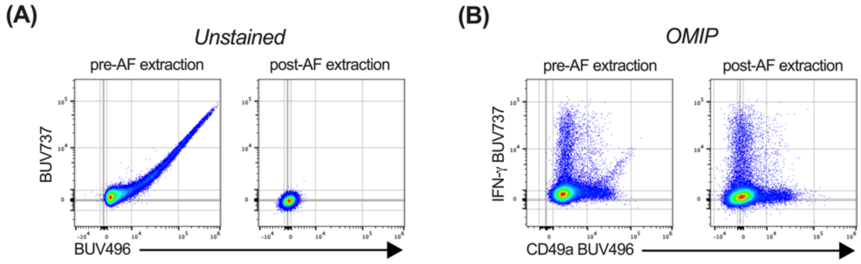


**Online Figure 6. Removal of cellular autofluorescence. (A)** Unstained lung single cells pre- and post-autofluorescence (AF) extraction demonstrating the presence of an AF streak within the UV7 channel, resulting in a false-positive BUV496 signal. **(B)** AF can be observed within the UV7 channel on viable CD45⁺ leukocytes prior to AF extraction, giving a false-positive IFN-γ⁺CD49a⁺ signal. AF extraction results in the successful removal of this false-positive signal. Single cell suspensions were stimulated *ex vivo* with PMA, Ionomycin, Brefeldin A, IL-12 and IL-18 (pro-ILC1 stimulation cocktail) for 5 hours prior to staining.

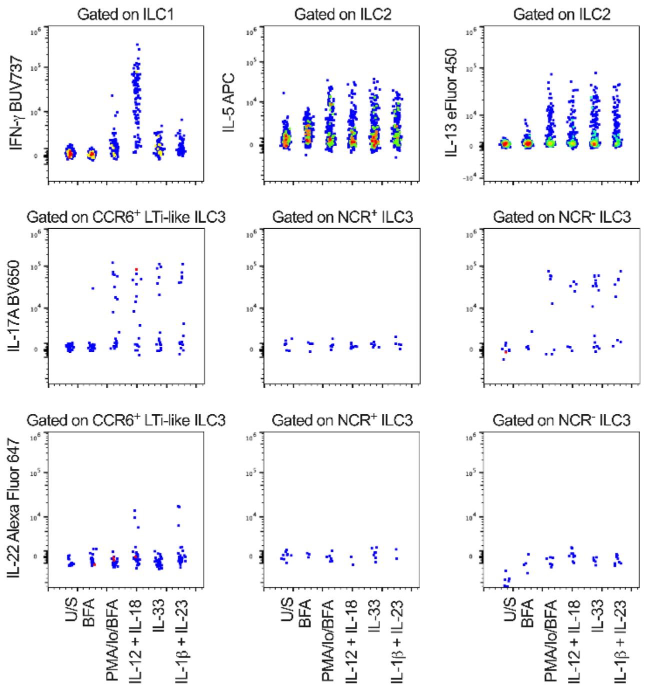


**Online Figure 7. Lung ILC subset cytokine expression in response to pro-ILC activating cytokines.** Comparison of ILC subset-specific cytokine expression within the lung following 5-hour *ex vivo* culture in R10F alone (U/S: unstimulated), R10F containing BFA only, R10F containing PMA/Ionomycin/BFA stimulation cocktail only, or R10F containing PMA/Ionomycin/BFA with either IL-12 + IL-18, IL-33 or IL-1β + IL-23.

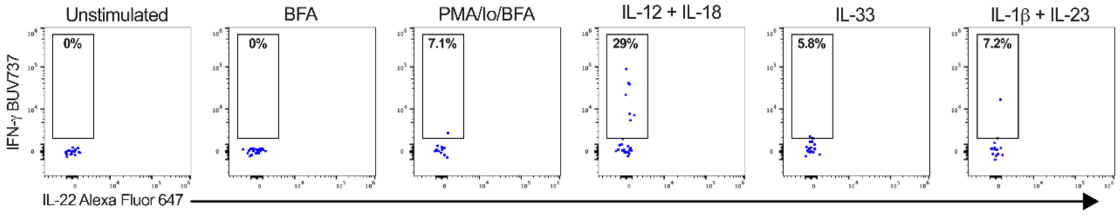


**Online Figure 8. Lung NCR⁺ ILC3 IFN-γ expression in response to pro-ILC activating cytokines.** Representative plots demonstrating IFN-γ expression by lung NCR⁺ ILC3s following 5-hour *ex vivo* culture in R10F alone (unstimulated), R10F containing BFA only, R10F containing PMA/Ionomycin/BFA stimulation cocktail only, or R10F containing PMA/Ionomycin/BFA with either IL-12 + IL-18, IL-33 or IL-1β + IL-23.

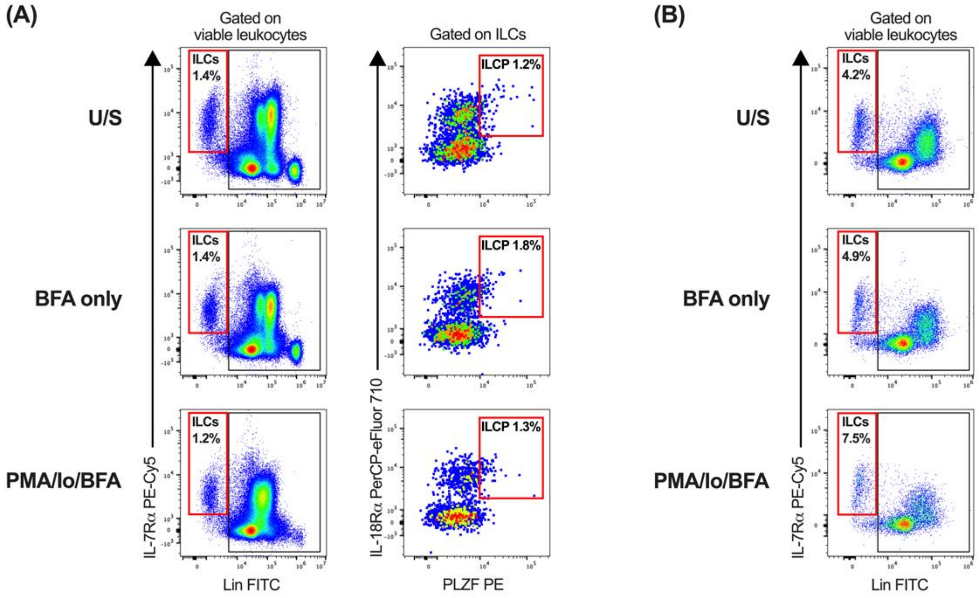


**Online Figure 9. ILC gating strategy validation in the absence of pro-ILC activating cytokines.** Comparison of staining profiles for ILCs and ILCPs in the lung **(A)** and ILCs in the siLP **(B)** following 5-hour *ex vivo* culture in R10F alone (U/S: unstimulated), R10F containing BFA only or R10F containing PMA/Ionomycin/BFA stimulation cocktail. Data are pre-gated to remove cellular debris (SSC v FSC), RBC contamination (SSC-A v SSC-B-A) and doublets (FSC-A v FSC-H), followed by selection of viable (LIVE/DEAD™ Blue$^{-}$) CD45$^{+}$ leukocytes.

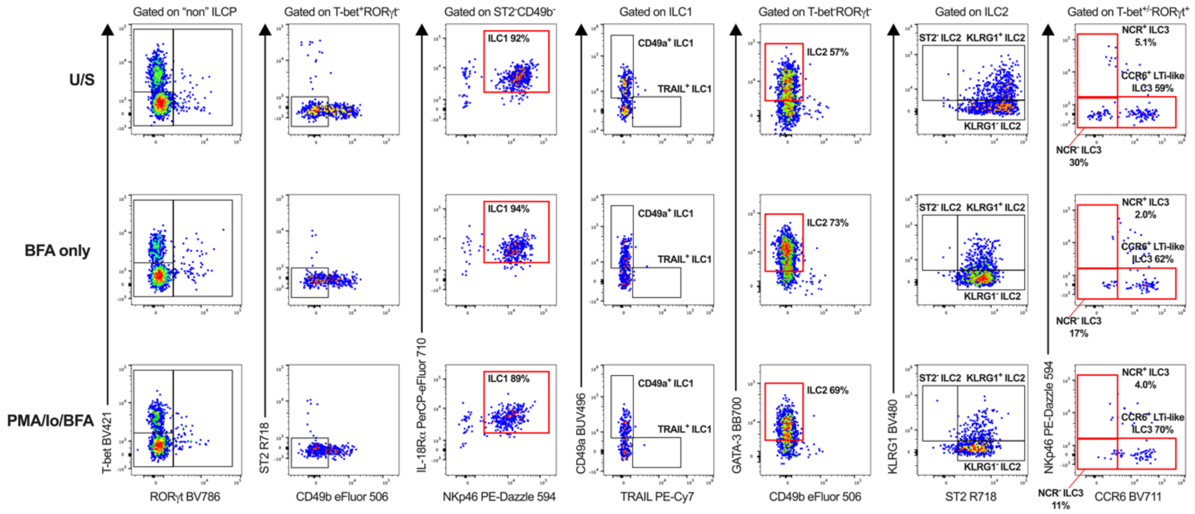


**Online Figure 10. Lung ILC subset gating strategy validation in the absence of pro-ILC activating cytokines.** Comparison of ILC subset staining profiles within the lung following 5-hour *ex vivo* culture in R10F alone (U/S: unstimulated), R10F containing BFA only or R10F containing PMA/Ionomycin/BFA stimulation cocktail. Data are pre-gated remove cellular debris (SSC v FSC), RBC contamination (SSC-A v SSC-B-A) and doublets (FSC-A v FSC-H), followed by selection of viable (LIVE/DEAD™ Blue$^{-}$) CD45$^{+}$ leukocytes.

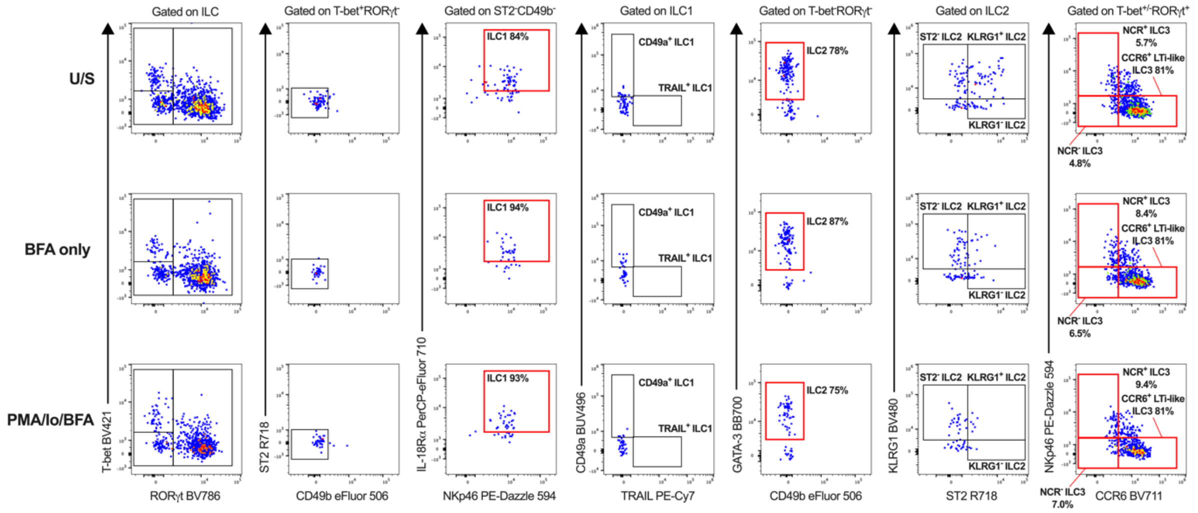


**Online Figure 11. siLP ILC subset gating strategy validation in the absence of pro-ILC activating cytokines.** Comparison of ILC subset staining profiles within the siLP following 5-hour *ex vivo* culture in R10F alone (U/S: unstimulated), R10F containing BFA only or R10F containing PMA/Ionomycin/BFA stimulation cocktail. Data are pre-gated remove cellular debris (SSC v FSC), RBC contamination (SSC-A v SSC-B-A) and doublets (FSC-A v FSC-H), followed by selection of viable (LIVE/DEAD™ Blue⁻) CD45⁺ leukocytes.

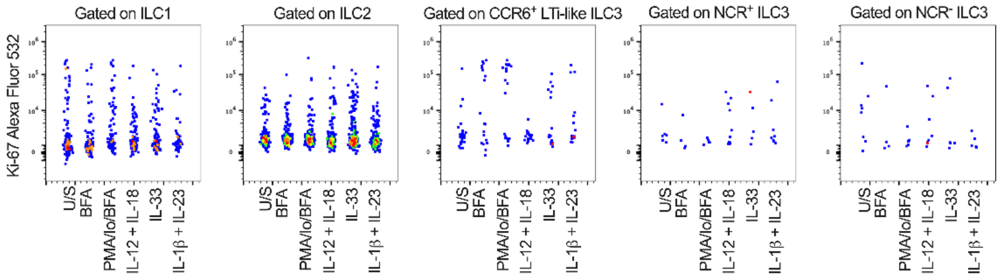


**Online Figure 12. Lung ILC subset Ki-67 expression in response to pro-ILC activating cytokines.** Comparison of ILC subset-specific Ki-67 expression within the lung following 5-hour *ex vivo* culture in R10F alone (U/S: unstimulated), R10F containing BFA only, R10F containing PMA/Ionomycin/BFA stimulation cocktail only, or R10F containing PMA/Ionomycin/BFA with either IL-12 + IL-18, IL-33 or IL-1β + IL-23.

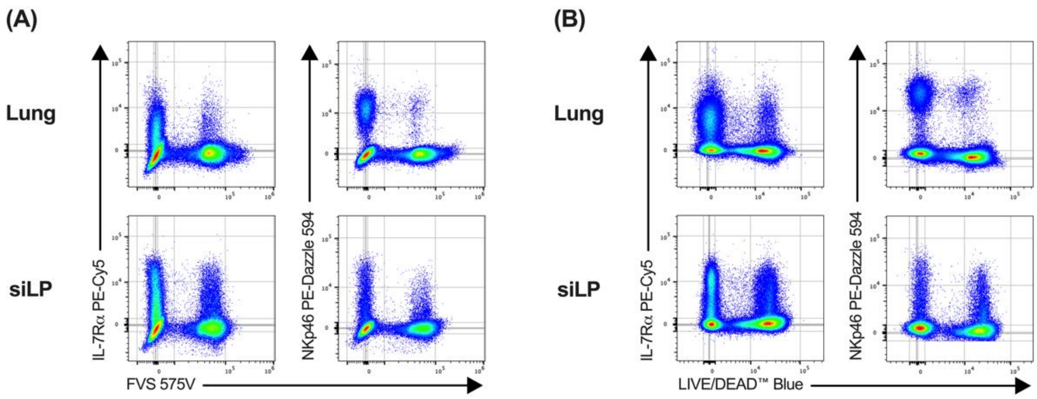


**Online Figure 13. Viability stain fluorochrome iterations during panel optimisation.** Non-viable cells were initially stained with FVS 575V in the original panel **(A)**, prior to replacement with LIVE/DEAD™ Blue in the final iteration of the OMIP **(B)** to reduce issues associated with spreading as identified in the SSM. Data are lung and siLP pre-gated to remove cellular debris (SSC v FSC) , RBC contamination (SSC-A v SSC-B-A) and doublets (FSC-A v FSC-H).

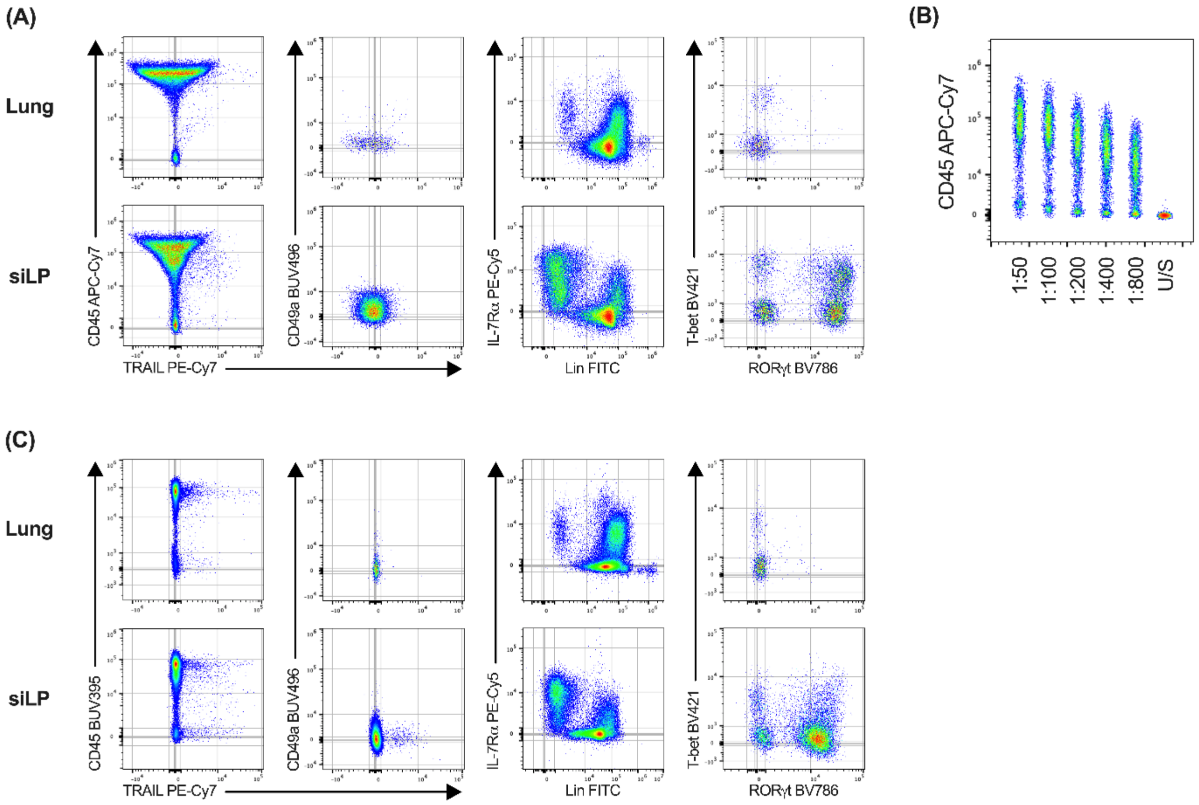


**Online Figure 14. CD45 fluorochrome iterations during panel optimisation.** Comparison of staining profiles in the lung and siLP using CD45 conjugated to APC-Cy7 **(A)** during the initial panel design and its associated titration **(B)**, and BUV395 **(C)** in the final OMIP. Single cell suspensions were stimulated *ex vivo* with PMA, Ionomycin and Brefeldin A (base stimulation cocktail) for 5 hours prior to staining. Data are pre-gated to remove cellular debris (SSC v FSC), RBC contamination (SSC-A v SSC-B-A), doublets (FSC-A v FSC-H) and non-viable (LIVE/DEAD™ Blue$^+$) cells.

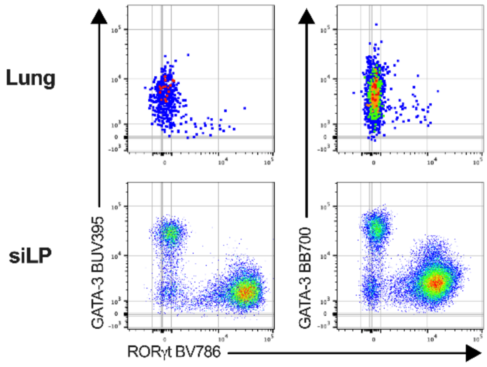


**Online Figure 15. GATA-3 fluorochrome iterations during panel optimisation.** Cells were initially stained with GATA-3 BUV395 for the characterisation of ILC2 in the original panel, prior to replacement with GATA-3 BB700 in the final iteration of the OMIP to improve positive signal resolution. Single cell suspensions were stimulated *ex vivo* with PMA, Ionomycin and Brefeldin A for 5 hours prior to staining. Data are pre-gated on lung and siLP IL-7Rα$^+$Lin$^-$ ILCs.

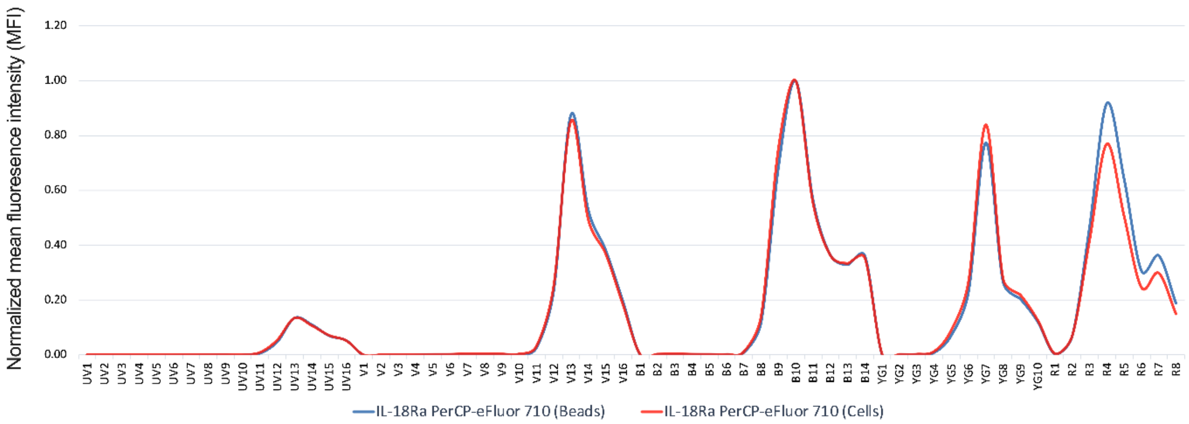


**Online Figure 16. Reference control optimisation.** Representative plot demonstrating potential inaccuracies when comparing cell- and bead-based reference controls. Significant deviation from the expected spectral profile of IL-18Rα PerCP-eFluor 710 was observed within the YG7, R4 and R7 channels when comparing stained lung single cell suspensions (red) to UltraComp™ eBeads (blue). As a results, a cell-based reference control is the only option for accurate unmixing of this fluorochrome.

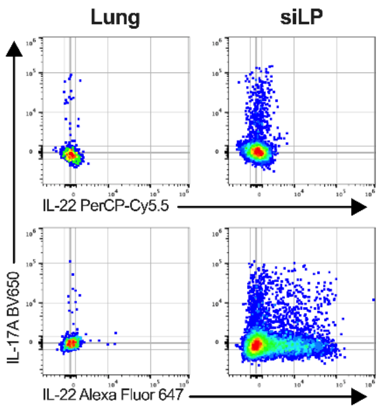


**Online Figure 17. Initial panel design using IL-22 PerCP-Cy5.5.** Cells were initially stained with IL-22 PerCP-Cy5.5 in the original panel, prior to replacement with IL-22 Alexa Fluor 647 in the final iteration of the OMIP to improve reference control quality and positive signal resolution. Single cell suspensions were stimulated *ex vivo* with PMA, Ionomycin, Brefeldin A, IL-1β and IL-23 (pro-ILC3 stimulation cocktail) for 5 hours prior to staining. Data are pre-gated on lung and siLP IL-7Rα$^+$Lin$^-$ ILCs.

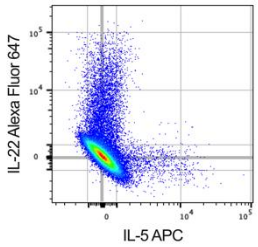


**Online Figure 18. Combination of Alexa Fluor 647 and APC.** This combination of highly overlapping fluorochromes in the same panel is achievable when the associated markers are not co-expressed. Single cell suspensions were stimulated *ex vivo* with PMA, Ionomycin, Brefeldin A, IL-1β and IL-23 (pro-ILC3 stimulation cocktail) for 5 hours prior to staining. Data are siLP pre-gated to remove cellular debris (SSC v FSC) , RBC contamination (SSC-A v SSC-B-A) and doublets (FSC-A v FSC-H), followed by selection of viable (LIVE/DEAD™ Blue$^-$) CD45$^+$ leukocytes.

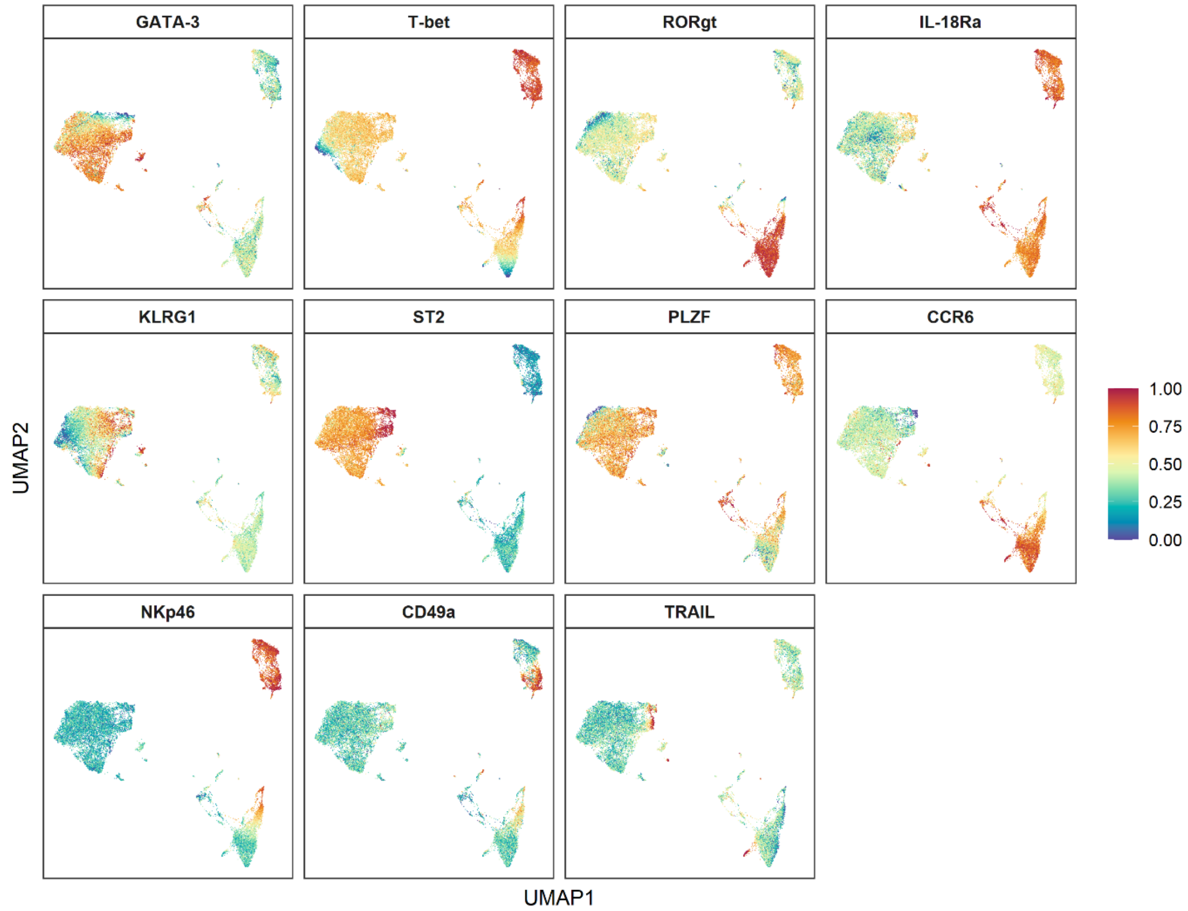


**Online Figure 19. Distribution of phenotypic markers on mouse ILCs.** Heatmap visualisation of phenotypic marker distribution across each of the distinct ILC UMAP clusters. Dimensionality reduction and UMAP was performed on 3,600 IL-7R$\alpha^{+}$Lin$^{-}$CD49b$^{-}$ ILCs from n=18 lung (n=3 unstimulated; n=3 BFA only; n=3 PMA/Io/BFA; n=3 IL-12 + IL-18; n=3 IL-33; n=3 IL-1$\beta$ + IL-23) and n=18 siLP (n=3 unstimulated; n=3 BFA only; n=3 PMA/Io/BFA; n=3 IL-12 + IL-18; n=3 IL-33; n=3 IL-1$\beta$ + IL-23) samples concatenated prior to unbiased clustering. Colours depict scaled marker expression from 0 – 1.

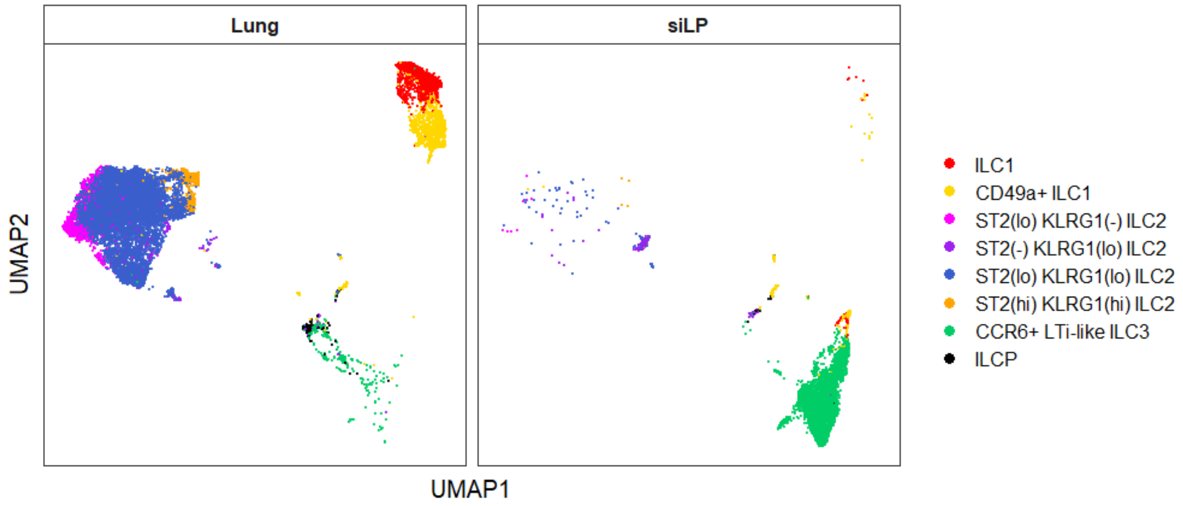

**Online Figure 20. Unsupervised clustering of mouse lung and siLP ILCs.** High-dimensional analysis of ILC subset abundance stratified on tissue localisation recapitulates the population dynamics observed following manual gating. Dimensionality reduction and UMAP was performed on 3,600 IL-7Rα$^+$Lin$^-$CD49b$^-$ ILCs from n=18 lung (n=3 unstimulated; n=3 BFA only; n=3 PMA/Io/BFA; n=3 IL-12 + IL-18; n=3 IL-33; n=3 IL-1β + IL-23) and n=18 siLP (n=3 unstimulated; n=3 BFA only; n=3 PMA/Io/BFA; n=3 IL-12 + IL-18; n=3 IL-33; n=3 IL-1β + IL-23) samples concatenated prior to unbiased clustering, followed by tissue stratification.

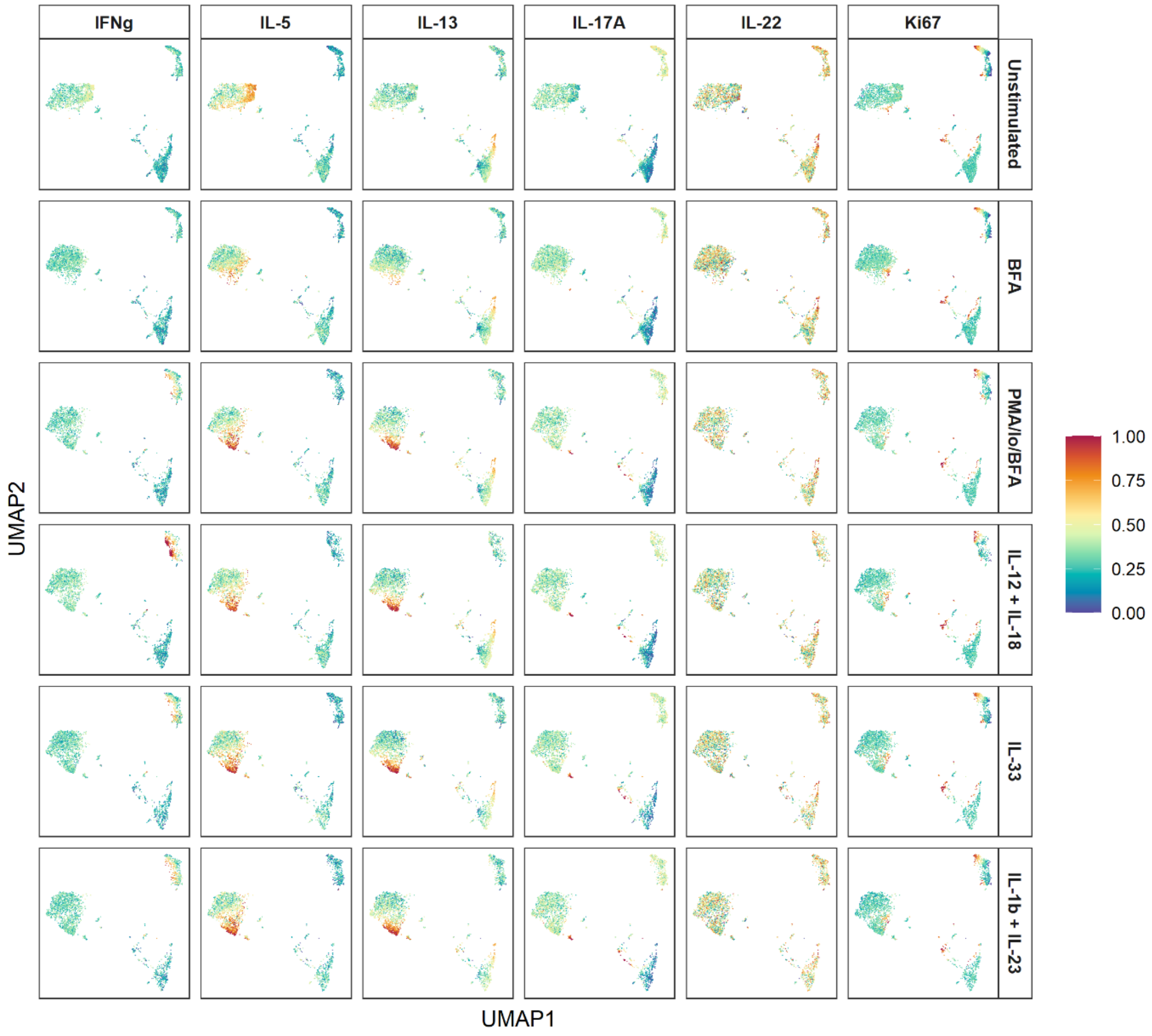


**Online Figure 21. Cytokine profiles and proliferative state of ILC subsets.** Heatmap visualisation of functional marker distribution across each of the distinct ILC UMAP clusters stratified on *ex vivo* stimulation condition. Dimensionality reduction and UMAP was performed on 3,600 IL-7Rα$^+$Lin$^-$CD49b$^-$ ILCs from n=18 (n=3 unstimulated; n=3 BFA only; n=3 PMA/Io/BFA; n=3 IL-12 + IL-18; n=3 IL-33; n=3 IL-1β + IL-23) lung and n=18 siLP (n=3 unstimulated; n=3 BFA only; n=3 PMA/Io/BFA; n=3 IL-12 + IL-18; n=3 IL-33; n=3 IL-1β + IL-23) samples concatenated prior to unbiased clustering. Colours depict scaled marker expression from 0 – 1.
